# Establishing *Ac*/*Ds* Starter Lines for Large-Scale Transposon-Tagged Mutagenesis in Tomato

**DOI:** 10.1101/2023.07.31.550784

**Authors:** Alka Kumari, Rachana Ponukumatla, Arun Kumar Pandey, Yellamaraju Sreelakshmi, Rameshwar Sharma

## Abstract

Tomato (*Solanum lycopersicum*), a model system for ripening of fleshy fruits, has ∼40,000 genes predicted by *in silico* homology-based annotation. However, functional validation is lacking for most annotated tomato genes. Among the strategies for functional annotations, transposon-tagged mutagenesis is the most powerful approach. Transposon-tagged genes can be functionally validated by phenotyping and activation tagging. However, the lack of a robust *in planta* transformation system precludes large-scale transposon-mutagenesis of tomato. To overcome this limitation, we developed two sets of starter lines in tomato, each carrying maize transposon element (*Ds*) and transposase (*Ac*), respectively. The *Ds* and *Ac* lines were crossed to allow the *Ac-*mediated transposition of the *Ds* in the F_1_ generation. In the F_2_ generation, the location of excised *Ds* at new sites in the tomato genome was monitored. The *Ds* transposition was interspersed on different chromosomes of the tomato, indicating unlinked transposition of the *Ds*. The analysis of DNA sequences flanking *Ds* showed random integration of *Ds* in intergenic, genes, and the promoter region of the genome. Our study paves the way for the generation of large-scale transposon-tagged tomato lines using *Ac*/*Ds* starter lines and provides a potential tool for the functional validation of genes in tomato.

**Short summary:** We developed two sets of starter lines in tomato, carrying maize transposon element (*Ds*) and transposase (*Ac*), to enable large-scale transposon-mutagenesis, facilitating functional validation of tomato genes and for generating an insertional mutant resource in tomato.

## Introduction

The impending climate changes and burgeoning human population have placed a marked emphasis on doubling global food production by 2050. Parallel to cereals that provide the most calorific value, there is an impetus to increase the yield and nutraceuticals in vegetable crops. Tomato, an important globally grown crop, is enriched in several nutraceuticals bereft in cereals. Like other crops, domestication pressure in tomato led to a decline in nutritional value, flavor, and disease resistance due to genetic erosion (**Bauchet and Causse, 2012; Tieman et al., 2017**). Genome resequencing of a large number of tomato cultivars, along with several wild relatives, has highlighted the extent of genetic diversity loss in modern tomato cultivars during domestication (**Aflitos et al., 2014; Gupta et al., 2020**).

The genetic diversity of a domesticated crop can be augmented by the introgression of chromosomal segments from wild relatives or by *de novo* diversity induction through induced mutagenesis (**Kulus, 2018**). Before the advent of the genomics era, the diversity induced by mutagenesis was harnessed by the visual selection of mutants displaying desired traits and backcrossing to the parent. The availability of genome sequences of crop plants expanded the scope of introgression of induced mutations by providing molecular markers (**Simko et al., 2021**). It also facilitated functional genomic analysis of genes mutated by chemical/radiation mutagens, T-DNA, or transposon insertion.

Among the various mutation protocols available, transposons occupy a prime position, as they allow the generation of a large collection of gene knockouts. The potential of autonomous transposable element activator (*Ac*) from maize, discovered by **McClintock (1951),** and its nonautonomous derivative dissociation (Ds) for gene tagging and insertion mutagenesis is well recognized (**Walbot, 1992**). The *Ac/Ds* elements also function in heterologous systems, as the *Ac*-encoded transposase protein is enzymatically active for the transposition of self or nonautonomous *Ds* elements.

Compared to T-DNA mutagenesis, the *Ac*/*Ds* transposon-based approach has several advantages. First, unlike the T-DNA mutagenesis system, which necessitates large-scale transformations, only a few primary transformants are needed to generate a large number of transposon-tagged lines. Second, *Ds* insertions can occur in somatic tissues or during gametogenesis. While the former is useful for activation tagging, insertion during gametogenesis results in *Ds* insertion at novel sites. Third, *Ds* is a nonautonomous element, and stable *Ds* mutants can be generated as *Ac* segregates away in later generations.

The maize *Ac*/*Ds* system has been successfully applied in several plant species for gene-tagging and insertion mutagenesis, such as Arabidopsis (**Parinov et al., 1999; Raina et al., 2002**), rice (**Greco et al., 2003; Kolensik et al., 2004**) and tomato (**Meissner et al., 2000; Carter et al., 2013**). In tomato, using *Ac*/*Ds* elements, the *CF-9* gene involved in plant resistance to the fungal pathogen *Cladosporium fulvum* was identified (**Jones et al., 1994**). Additionally, the *DCL* gene, required for chloroplast development and palisade cell morphogenesis, and the *FEEBLY* gene, involved in plant development, were also identified using *Ac*/*Ds* for gene tagging (**Keddie et al., 1996; van der Biezen et al., 1996**).

Two main strategies have been employed to generate *Ac*/*Ds*-based insertion mutation lines. The first approach involves placing the *Ac* and *Ds* elements in the same construct (Greco et al., 2003), resulting in transposed plants without the need for crossing. However, to segregate Ds-only plants, a complex screening protocol is needed. This strategy was applied by **Meissner et al. (2000)** and **Carter et al. (2013)** to generate *Ds*-tagged lines in tomato. The second strategy entails placing the *Ac* and *Ds* elements on separate constructs and independently transforming them into starter lines (**Kolesnik et al., 2004**). The Ds element can then be excised from its original site and transposed to new locations through the crossing of *Ac* and *Ds* plants, allowing for the selection of *Ds*-only plants.

In this study, we generated an insertional mutant population in tomato using the second strategy wherein we developed two independent sets of starter lines, one carrying the *Ac*- transposase and the other containing the *Ds*-transposon. Ac-transposase was detected using GFP as a reporter gene, and *Ds*-transposon had both GFP and RFP as reporters. Upon mobilization to a new site, the *Ds* excised itself, carrying RFP along but leaving GFP at the original site. The selection of *Ds*-transposed plants involved a much simpler protocol where Ds-only plants were selected based on GFP-negative and RFP-positive plants. This two-starter line system is more flexible and reliable than a single transformed line for generating a large insertion mutagenized population.

## Results

For transposon mutagenesis of tomato, we selected a strategy involving two independent parental lines, one bearing *Ac*-transposase (*Ac*-**Activator**) and another bearing Ds- transposon (*Ds*- **Dissociation**). It was surmised that crossing these two parental lines would lead to excision and reinsertion of the Ds element in F_1_ progeny. The F_2_ progeny can be analyzed for plants having mobilized Ds element at a new genomic location sans *Ac* transposase. To facilitate screening at the seedling stage, we chose the reporter genes GFP **(green fluorescent protein)** and DsRed/RFP **(red fluorescent protein)**, which can be screened by examining fluorescence or PCR. **Figure S1** shows sequential steps for *Ac/Ds* mutagenesis of tomato using the above strategy.

### Modifications of *Ac* and *Ds* constructs and raising of *Ac/Ds* parental lines

The constructs utilized for transposon mutagenesis were initially developed for rice, a monocot plant (**Sundaresan et al., 1995, Figure 1A**). Modifications were made to adapt these constructs for efficient expression and transformation in tomato, a dicot plant (**Figure 1B**). The *Ac* construct contained an immobilized *Ac-TPase* gene, while the *Ds* construct (derived from the wild-type *Ac-TPase* element) consisted of a nonautonomous *Ds* element with 222 bp at the 3’ end and 1785 bp at the 5’ end (**Figure 1**). The construct included a 4X Enhancer (4X EN) sequence to enable activation tagging. The modifications involved replacing the maize ubiquitin promoter with the Arabidopsis ubiquitin promoter and substituting the hygromycin-resistance gene with a kanamycin resistance gene (**Figure S2-S3**). The modified *Ac* construct included the *GFP* gene as a reporter, while the modified *Ds* construct included both the *GFP* and *RFP* genes as reporters (**Figure S4-5**). The modified *Ac* and *Ds* vectors were introduced into tomato cv. Arka Vikas through Agrobacterium-mediated genetic transformation. In the T_0_ generation, twenty-four *Ac* lines and twenty *Ds* lines were selected. Before initiating genetic crossing, the parental lines were stabilized up to the T_4_ generation for *Ac* and the T_2_ generation for *Ds*.

**Figure 1.**
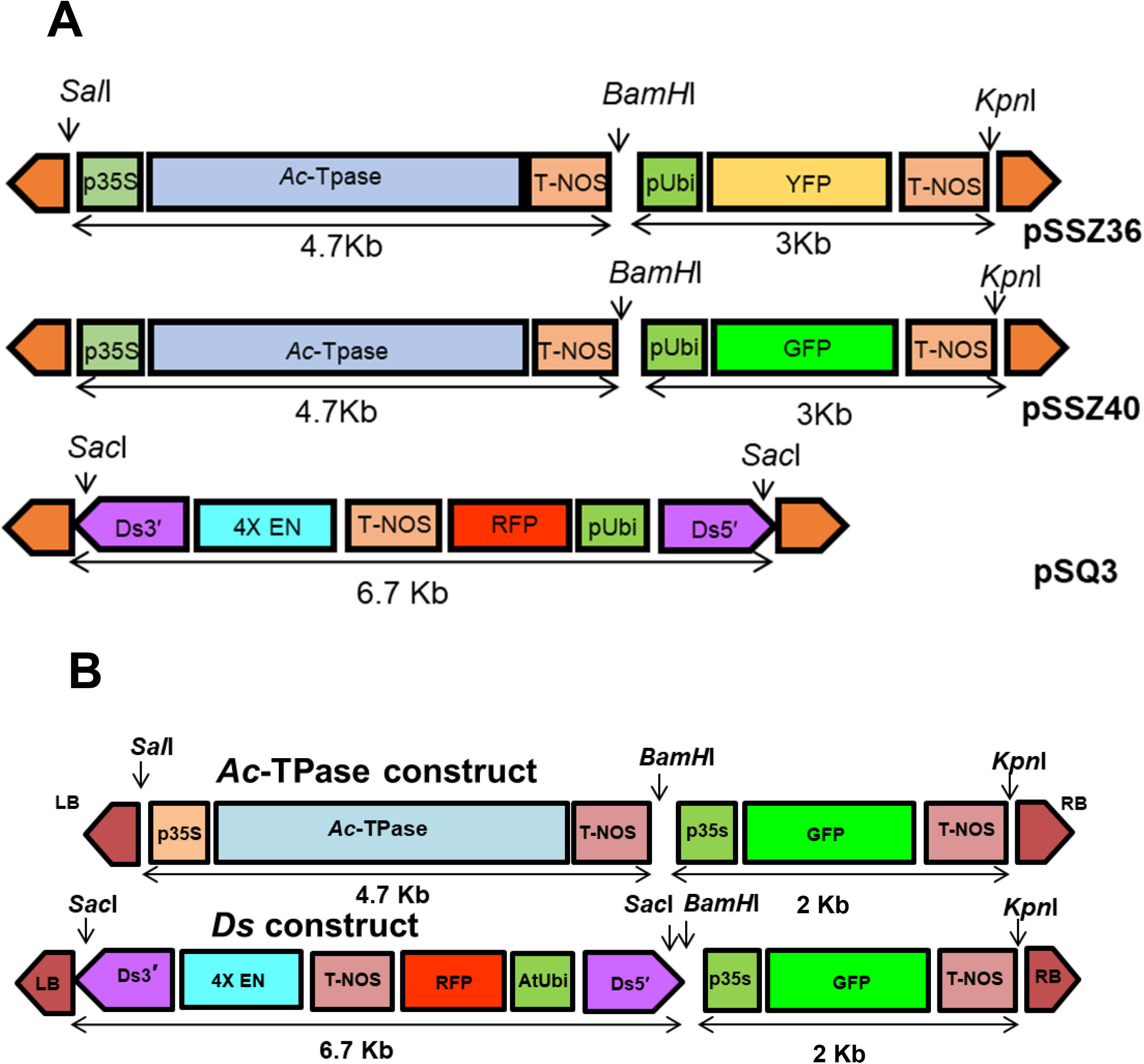
Constructs used for raising transposon-tagged lines in tomato. **A.** The *Ac* and *Ds* constructs used for insertional mutagenesis in rice. **B.** The modified *Ac* and *Ds* constructs used for insertional mutagenesis in tomato.

### Reporter gene expression in *Ac* and *Ds* lines

We screened the *Ac* and *Ds* lines using a kanamycin resistance assay, PCR amplification, and GFP/RFP fluorescence visualization (**Figure 2**). Based on the appearance/lack of bleaching, the plants were categorized as resistant or sensitive to kanamycin (**Figure S6, Table S1**). The transgenicity of kanamycin-resistant lines was also reconfirmed by PCR amplification of transgenes (**Figure S7**). Additionally, the integration of the transgene was also verified by Southern blotting (**Figure S8**). The possibility of Agrobacterium persistence was eliminated by multiplex PCR using the chromosomal virulence (*chv*) gene as an Agrobacterium marker, lycopene β-cyclase (*CYCB*), a single copy native gene of tomato, as a host plant marker, and *NPTII* as a transgene construct marker (**Figure S9**).

**Figure 2.**
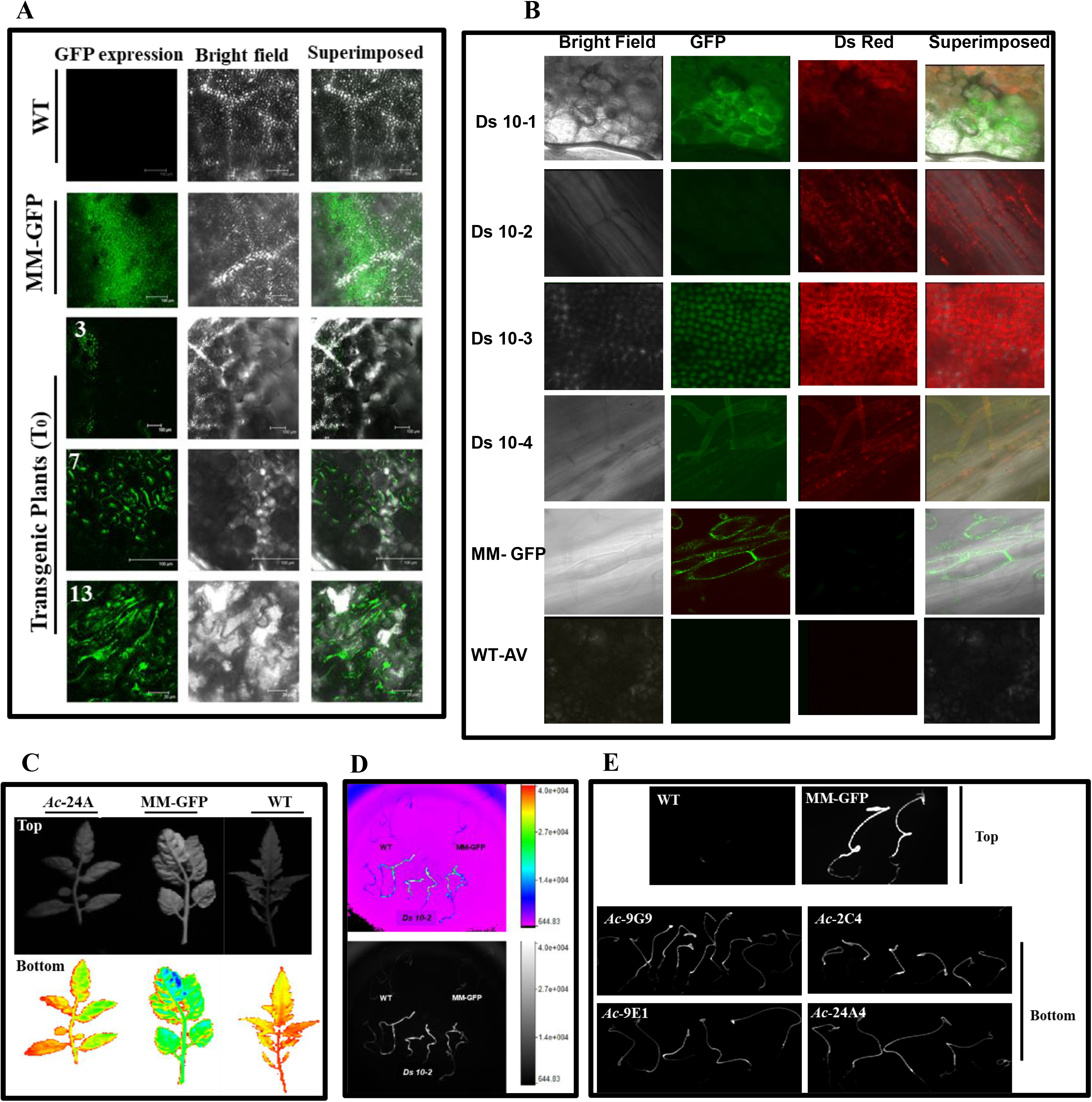
Imaging of GFP and RFP fluorescence in *Ac-* and *Ds* plants. WT (Arka Vikas) and transgenic GFP plants in the Moneymaker background were used as negative and positive controls, respectively. The emission of GFP and *RFP* fluorescence from the respective plants indicates the presence and expression of the *GFP* and *RFP* **transgenes. A-B.** GFP (**A**) and DsRed (RFP) (**B**) expression analysis in T_0_ *Ac* and *Ds* lines using confocal laser scanning microscopy. **C**. The top panel shows the pictures taken using the GFP filter, and the lower panel shows the photographs of the same leaves. The blue color in the leaf shows the saturation of *GFP* fluorescence. *Ac-*24A T_2_ is a Southern positive line. **D.** The top panel shows pictures of WT (left) and MM-GFP (right) seedlings taken with a *RFP* filter, and the lower panel shows RFP fluorescence from Southern-positive *Ds10-2* T_1_ transgenic line seedlings. **E.** The top panel shows pictures of WT (left) and MM-GFP (right) seedlings taken with a *GFP* filter. The bottom panel shows *GFP* fluorescence from the seedlings of *Ac* T_3_ lines. The pictures for **C-E** were taken in the Kodak imaging station.

In T_0_, *Ac* plants displayed distinct green fluorescence; *Ds* plants showed both green and red fluorescence, whereas untransformed plants exhibited no fluorescence (**Figure 2A-B**). The *Ds* T_0_ plants displayed GFP and RFP fluorescence in the leaves (**Figure 2B**). The expression of GFP in *Ac* lines was further confirmed in the T_2_ (leaves **Figure 2C**) and T_3_ (seedlings) generations (**Figure 2E**). The dark-grown T_2_ *Ds* seedlings (**Figure 2D**) displayed RFP fluorescence. However, the intensity of GFP and RFP fluorescence varied across different *Ac* and *Ds* lines, with some lines exhibiting weaker fluorescence. We also screened the F_2_ population using PCR amplification of the GFP and RFP genes to mitigate the risk of missing a *Ds*-mobilized line due to weak fluorescence. We selected seven *Ds* lines and four *Ac* lines for further analysis based on Southern blotting, fluorescence visualization, and PCR amplification assays.

### Generation of Ds-tagged population

To generate the F_1_ population, different combinations of *Ac* (Female ♀) and *Ds* (Male ♂) starter lines, each containing a single transgene copy, were manually crossed. Two independent sets of crosses were made. The first set of cross viz, *Ds-*2 X *Ac*-9E1, *Ds-*9 X *Ac*- 9E1, *Ds-*6 X *Ac*-9E1, *Ds-*8 X *Ac*-24A3, *Ds-*3 X *Ac*-2C4, *Ds-*1 X *Ac-*9G9, was analyzed for flanking sequence using FPNI-PCR. The second set of crosses with viz. *Ds-*10-2 X *Ac-*24-A, *Ds-*10-8 X *Ac-*9G, and *Ds-*10-1 X *Ac-*9G were analyzed using inverse PCR. The launch sites for *Ds-2*, *Ds-4,* and *Ds10-2* are given in **Table S2**. In either set of crosses, the presence of *GFP* and *RFP* fluorescence in F_1_ seedlings signified the successful transfer of *Ds* transgene (♂) (RFP^+^/GFP^+^) to the *Ac* (♀) (GFP^+^) lines, as maternal *Ac (♀)* lines were devoid of RFP^-^ (**Figure 3**). The presence of the *GFP* and *RFP* genes was further confirmed using PCR with transgene-specific primers. The F_1_ plants were allowed to self-pollinate, and the F_2_ plants were screened for *Ds* mobilization.

**Figure 3:**
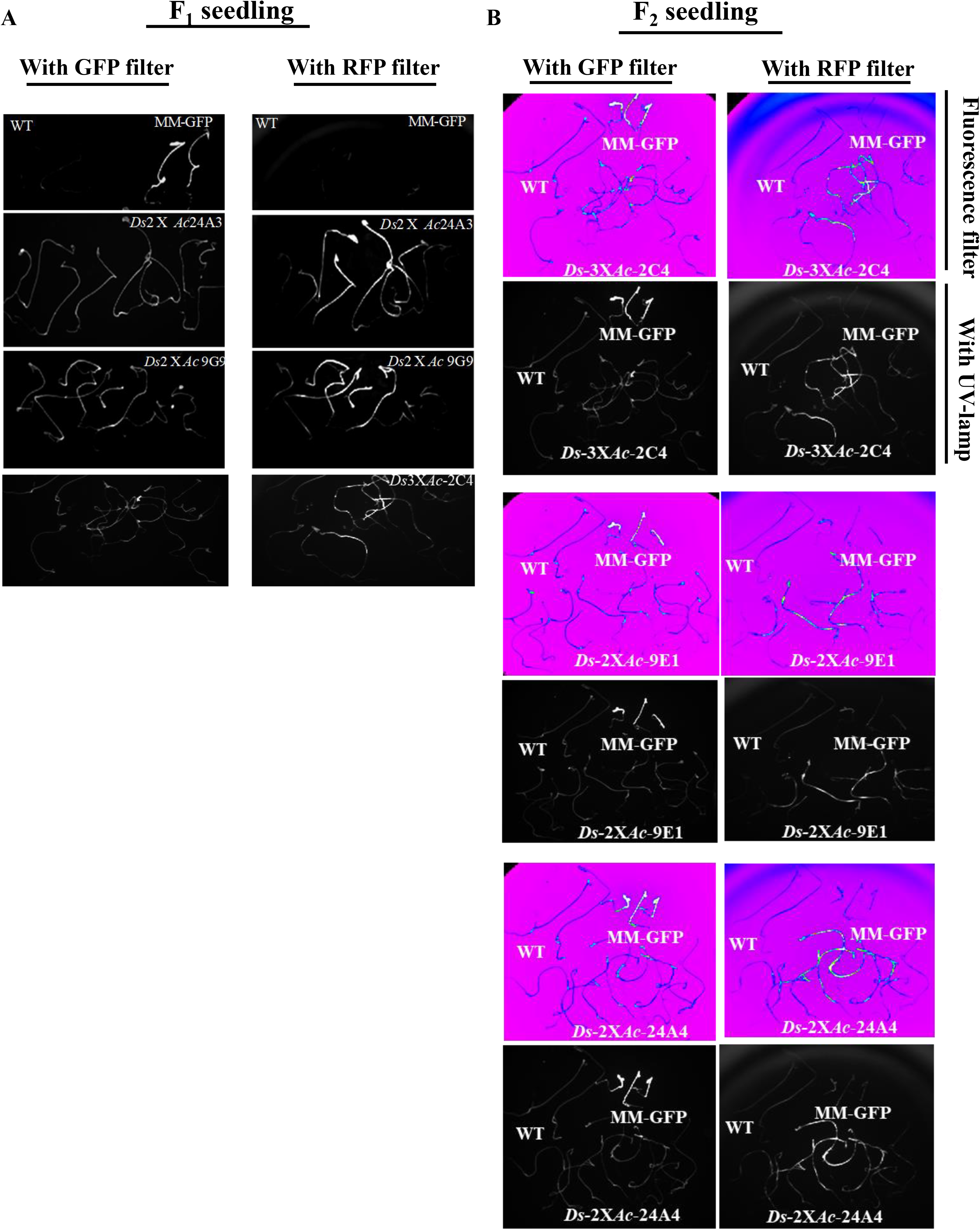
*In vivo* imaging of *RFP* and *GFP* fluorescence in F_1_ and F_2_ seedlings. **A**, Fluorescence of F_1_ seedlings, *Ds2* X *Ac-24A3*, *Ds2* X *Ac-9G9*, and *Ds*3 X *Ac*-*2C4* seedlings using GFP filter (left panel) and RFP filter (right panel). The seedlings were illuminated with a UV lamp. **B.** Image of F_2_ seedlings *Ds-2* X *Ac-24A4, Ds-3* X *Ac-2C4,* and *Ds-2* X *Ac-9E1* seedlings using GFP filter (left panel) and RFP filter (right panel). The seedlings were illuminated with a UV lamp and photographed using GFP and RFP filters. Untransformed seedlings (WT) do not show *GFP* and *RFP* fluorescence. The MM-GFP (positive control) lines show only *GFP* fluorescence but do not show *RFP* fluorescence.

For F_2_ plants, screening was performed to select GFP-negative (GFP^-^) and RFP-positive (RFP^+^) plants. The random phenotyping of a few F_2_ plants also showed morphological variations (**Figure S10**). Approximately 1000 F_2_ plants at the seedling stage were screened for *Ds* segregation for the second set of crosses. Out of 800 surviving F_2_ plants, 120 plants (15%) did not show amplification of the *GFP* gene by PCR, indicating the excision of the *Ds* element. These GFP^-^ plants were then screened using *RFP*-specific primers (**Figure 4**). Among these, sixty plants (7.5%) showed amplification of the *RFP* sequence (∼ 500 bp product). This confirmed the excision and reinsertion of the *Ds* elements. The genomic DNA from these GFP^-^/RFP*^+^* plants was further analyzed to determine the location of the mobilized *Ds* elements.

**Figure 4:**
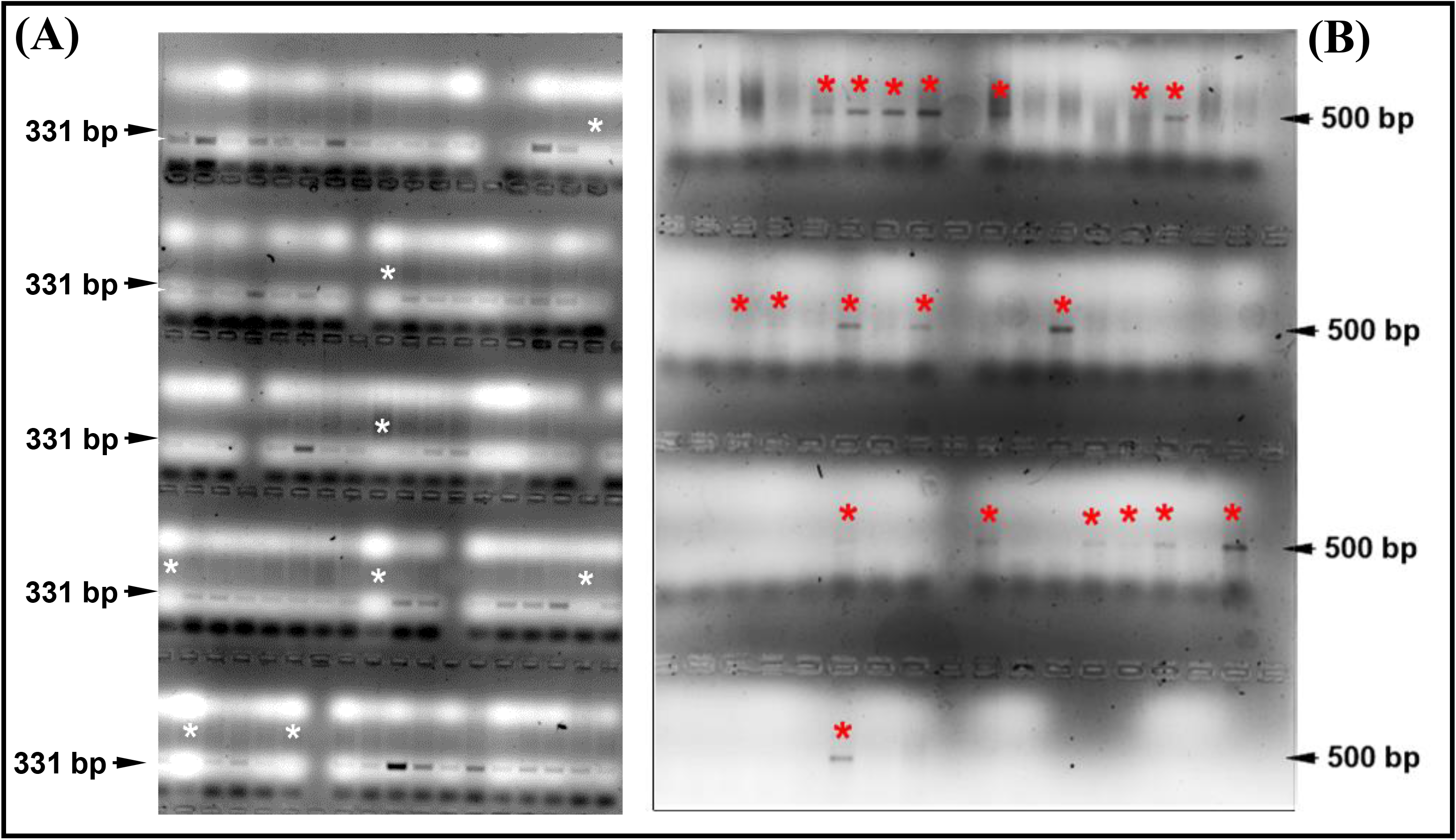
Analysis of *Ac-Ds* segregation in the F2 generation. **A.** PCR with *GFP-*specific primers. **Note:** A 331 bp amplicon indicates the presence of *Ac* and *Ds* or only *Ac*. A white asterisk indicates the absence of a *GFP* amplicon. **B.** PCR with *RFP-*specific primers. **Note:** A 500 bp amplicon indicates the presence of only mobilized *Ds* element. A red asterisk indicates that only the *RFP* amplicon was amplified.

### Isolation of Ds-Flanking Sequences from RFP^+^ Lines

The first set of crosses was analyzed using FPNI-PCR. However, this method could recover only ten transposition hits (**Dataset S1, #1-10**). Therefore, inverse PCR was used to identify the insertion sites of the transposed element in GFP^-^/RFP*^+^* plants for the second set of crosses. The amplified products were visualized using agarose gel electrophoresis and cloned into the pJET 2.0 vector to be sequenced (**Figure 5**). From the second set of RFP-positive plants, 46 flanking sequences were isolated, cloned, and sequenced (**Dataset S1, #11-56**). Among these 46 sequences, the longest amplicon was 1452 bp, and the average amplicon length was 1110 bp.

**Figure 5.**
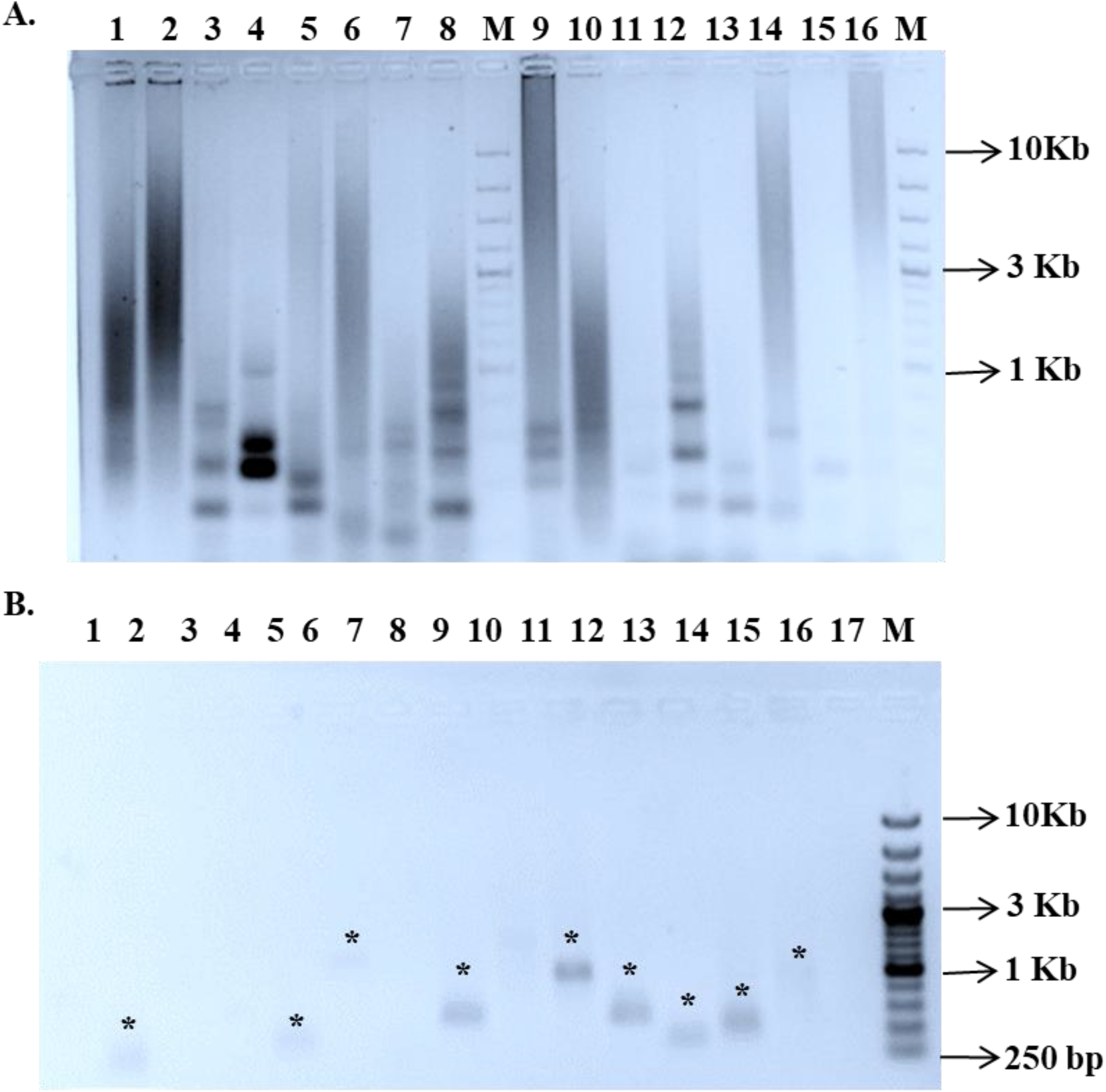
Inverse PCR to amplify *Ds*-tagged DNA from GFP^-^ and RFP^+^ *Ds* F_2_ lines. **A.** Products of inverse PCR analysis using DNA isolated from leaves of *Ds-*only plants. **B.** Gel-purified amplicons of flanking sequences obtained after PCR. Note: The asterisks show the amplicons of flanking sequences in different samples. **M:** 1 Kb Marker. Lanes 1-16, *Ds*-only plants.

Sequence homology studies using a BLAST search against the tomato genome showed that all flanking sequences had hits to the tomato genome but had only 24 unique hits (52%). The highest insertions were found on chromosome 3, followed by chromosomes 8 and 10, with six insertions each, and chromosome 2, with four insertions. Chromosome 4 has two insertions, with the remaining chromosomes at a single insertion, each mapped on them (**Table S3**). However, no insertion was obtained on chromosome 1 (**Dataset S1, Figure 6**). Twenty-one insertions were mapped to chromosome 3, of which 19 were at the same chromosomal location. All three parental *Ds* (10-2, 10-8, 10-1) lines had the insertion in chromosome 3 at the exact location (SL3.0ch03, 26676740- 26676787), as they were the progeny of a single T_0_ line number *Ds-*10.

**Figure 6.**
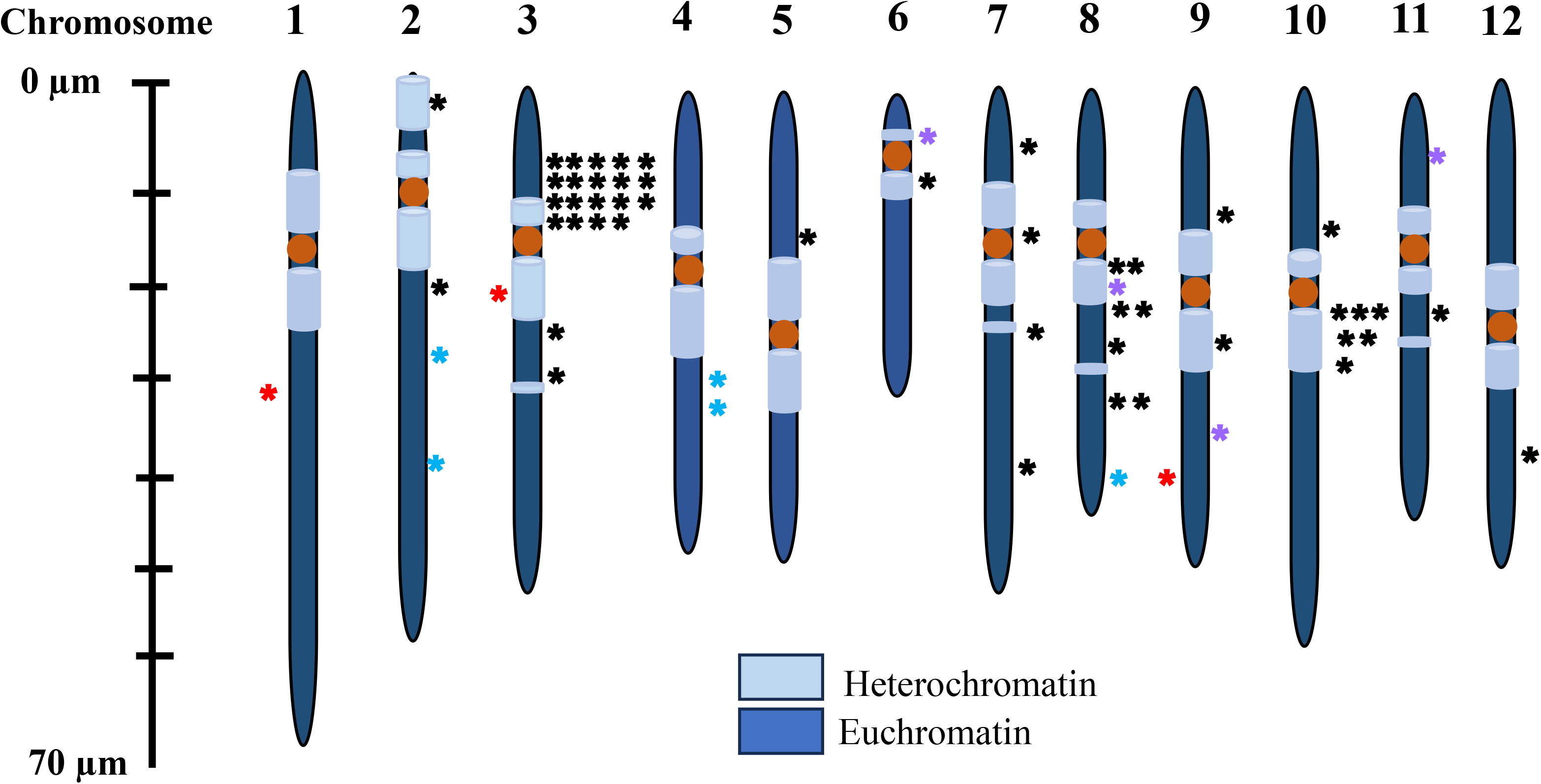
Schematic representation of the distribution of *Ds* on twelve chromosomes of tomato. Out of 12 chromosomes, transposon insertion was found on 11 chromosomes. The blue asterisks indicate insertions in genic regions, while the black asterisks indicate insertions in intergenic regions. The purple asterisk indicates the insertions whose launch site was not known. The red asterisks on the left side of chromosomes 1 (*Ds-2*), 3 (*Ds10-2*), and 9 (*Ds-8*) indicate the launch site. The data are from both sets of crosses. The coordinates of the insertion sites are given in Dataset S1.

### Distribution of transposon insertions within intergenic and intragenic regions

A total of 8.7% of insertions occurred in the intragenic region comprising exons, introns, untranslated regions (UTRs), and regions within 1 Kb upstream or downstream of a gene (exons, introns, plus UTRs). The intergenic regions had an insertion frequency of approximately 91%. The insertion frequency in CDS was also 2.7% (**Figure S11**). The frequency of insertions in the introns and promoters was similar to CDS, accounting for 2.7% each. The analysis of *Ds* insertions revealed that *Ds* inserted into the genic regions of the ABC transporter family protein, DNA repair endonuclease, RNA-binding (RRM/RBD/RNP motifs) family protein, and ATP synthase α subunit genes. Other insertions were present in the intergenic regions of genes such as eukaryotic aspartyl protease family protein and BED zinc finger, cytochrome P450 and phosphatidylinositol N-acetyl glucosamine transferase subunit P-like protein, xanthine/uracil permease family protein, small nuclear ribonucleoprotein, 2- oxoglutarate (2OG) and Fe(II)-dependent oxygenase superfamily protein, and tetratricopeptide repeat (TPR)-like superfamily protein. (**Database S1**). Almost 40% of the flanking sequences showed the same insertion site upstream of the auxin efflux carrier family protein and downstream of the clade XVI lectin receptor kinase.

The location of the *Ds* insertion site relative to the nearest upstream and downstream gene and the approximate GC percentage of the insertion site were also determined for the flanking sequences (**Dataset S1**). The approximate percentage of the GC content of the insertion sites in the tomato genome range was 30-100% (**Dataset S1**). The five genic insertions had a high percentage of GC content; insertions on chromosome 2 in RNA-binding (RRM/RBD/RNP motifs) family protein and plastid transcriptionally active 14 protein had GC contents of 100% and 57%, respectively. The two insertions on chromosome 4 in the ATP synthase alpha subunit and ABC family transporter protein had GC contents of 90% and 95.5%, respectively. The insertion on chromosome 8 in DNA repair endonuclease XPF had a GC content of 85%.

## Discussion

Considerable efforts have been directed to generate resources for functional genomics in tomato after the release of the complete tomato genome sequence (**The Tomato Genome Consortium, 2012**). Most resources were generated using EMS-mutagenized lines of tomato, and mutated genes were identified using classical TILLING (Targeting Induced Local Lesions in Genomes) (**Okabe et al., 2011; Sharma et al., 2021**), NGS-based TILLING (**Gupta et al., 2017**), and whole genome sequencing (**Shirasawa et al., 2016; Yano et al., 2019; Gupta et al., 2022**). However, EMS, being a chemical mutagen, generates a large number of mutations across the genome, necessitating extensive backcrossing for functional validation of an EMS mutant.

Compared to EMS, T-DNA mutagenesis allows targeted insertion of T-DNA into the genome. Moreover, the inserted T-DNA serves as a tag to identify the inserted loci and evaluate the associated phenotype. However, T-DNA mutagenesis relies on Agrobacterium-mediated transformation, which limits its application for large-scale mutagenesis, as tomato lacks a robust *in planta* transformation system. In tomato, **Pérez-Martín et al. (2017),** using 22,700 explants, generated 7,842 transgenic lines, of which 2,282 plants were tetraploid. The occurrence of a high number of tetraploids limits the utility of T-DNA tagging for functional genomics, as before the usage of each T-DNA-tagged line, its ploidy has to be ascertained. Moreover, a huge number of transformations are required to achieve whole genome saturation with T-DNA mutations.

We used maize activator/dissociation (*Ac*/*Ds*) transposons for the knockout of genes for insertional mutagenesis, which required a transformation step only to generate the *Ac* and *Ds* starter lines. A significant concern in the *Ac/Ds* transposition system is distinguishing *Ds-* only lines after *Ds* insertion at a new site. In rice, **Kolesnik et al. (2004)** identified *Ds-*only lines by first screening for GFP-negative plants and then subjecting them to the herbicide BASTA to identify BASTA-resistant seedlings bearing transposed *Ds* elements. Our strategy also consisted of two-level screening; the first was selecting GFP-negative plants, which eliminated both *Ac* starter lines and untransposed *Ds* plants. In the second step, the RFP- positive plants were selected to determine the new site of *Ds* transposition.

The robustness of the selection of *Ds* transposed to a novel site is determined by the extent of the expression of the linked markers. **Kolesnik et al. (2004)** reported that nearly 20% of GFP-negative plants in rice did not exhibit BASTA resistance. In plants, factors such as chromatin structural variation and neighboring genes within ≤ 20 Kb can repress or activate the transgene (**Akhtar et al., 2013**). Consistent with this, we observed variations in the fluorescence of GFP and RFP reporters in our study. To overcome this, we used PCR amplification of the reporter genes as the final step to identify the *Ds*-tagged lines at new insertion sites. The identification of new *Ds* transposition sites in our study shows the successful implementation of a two-component *Ac*/*Ds* system in tomato utilizing a negative selection of *Ac* transposase as well as of untransposed *Ds* using *GFP* as a reporter.

It is apt to compare our study with earlier studies where the *Ac*/*Ds* system was used to generate mutant lines in Micro-Tom (**Meissner et al., 2000**) and M82 (**Carter et al., 2013**). Although **Meissner et al. (2000)** caused 2932 F_3_ families, the identity of transposed loci could be determined only for 28 genes based on ESTs. **Meissner et al. (2000**) used a more complex screening protocol than our study for selecting F_2_ seedlings for germinally stable transposition. They used acetolactate synthase resistance as a marker for excision, hygromycin or kanamycin resistance for reinsertion, and naphthalene acetamide resistance for stabilization of transposition. Before this screening, the F_1_ plants were also screened for chlorosulfuron resistance to select plants containing *Ds* insertion. However, the potential use of the above population remained limited, as the tomato genome sequence became available only in 2012. A drawback of Micro-Tom is its miniature phenotype determined by a combination of hormonal and light-signaling mutations, which may obfuscate the expression of traits related to agronomic performance (**Carvalho et al., 2011**).

Using micropropagation, **Carter et al. (2013)** raised 25 T_1_ lines from a single T_0_ line and obtained 509 independent *Ds*-tagged lines. However, one major issue was the absence of GFP expression directed by a monocot promoter, which failed to work in dicot tomato. This limited the screening of transgenic plants based on a single hygromycin painting assay. In contrast, our starter lines do not suffer from these drawbacks, as these are raised in a commercial cultivar, and constructs are driven from dicot-specific promoters. Moreover, our lines can be used as a resource as a launch pad for generating mutants to study functional genomics and crop performance traits.

A major advantage of using a transposase-bearing line and a few transposon-carrying lines as starter lines is that one can generate numerous mutant lines by crossing these two lines. However, to generate mutants on a genome-wide scale, *Ds* should randomly mobilize across the different chromosomes of the tomato away from its launch site. It has been reported that *Ds* gives both linked and unlinked transpositions in tomato (**Osborne et al., 1991; Healy et al., 1993**). It contrasts with monocots, where *Ds* mostly gives rise to linked transpositions (**Smith et al., 1996**). In contrast to monocots, we observed both linked and unlinked transposition with respect to the launch site of *Ds10*-2 located on chromosome 3 of tomato.

An unusual aspect was multiple transpositions of the *Ds* at a single site on chromosome 3. In Arabidopsis, **Parinov et al. (1999),** using the *Ac*/*Ds* element, reported the existence of two transposition hot spots adjacent to nucleolus organizer regions NOR2 and NOR4. Similarly, in rice, the distribution of transposons on different chromosomes revealed a preferential transposition of *Ds* into a hot spot on chromosome 7 (**Kolensik et al., 2004**). In tomato, too, there is a preferential transposition of *Ds* on the same chromosome as the launch pad (**Bríza et al., 2002**). In the classical view, nineteen insertions at the same site may represent a hot spot for transposon insertion. It is also plausible that these 19 plants are siblings from a single F_1_ parent, where transposition occurred before gametogenesis. Nonetheless, more detailed studies are needed to confirm this possibility. It is more likely that the *Ds* 10-2 launch pad is in the vicinity of the above multiple insertion site; the *Ds* element has preferentially transposed to this site, giving rise to linked transpositions. These insertions on the same chromosome may arise due to chromatin accessibility to the transposase enzyme in the vicinity of the launch site.

Except for chromosome 1, the transposons were randomly dispersed on the other ten chromosomes of tomato, highlighting many unlinked transpositions. As RFP-only lines were examined, the number of observed unlinked transpositions might likely be lower than the actual ones. We might have missed those F_2_ lines where *Ds* transposed to a new location, but either empty *Ds* launchpad or *Ac* transposes bearing GFP did not segregate from *Ds* transposed to a new site. Nevertheless, the distribution of linked and unlinked transposition in our study is consistent with earlier observations that, unlike monocots, in tomato, both linked and unlinked transpositions occur at nearly the same frequency (**Osborne et al., 1991)**. Since there are no specific hot spots for unlinked transpositions, using a few *Ds* starter lines with a single *Ac* transposase line may lead to genome-wide *Ds* insertions.

It is believed that the distribution of linked versus unlinked transposition is determined by the developmental stage of the F_1_ plant at which the transposition takes place. Transposition occurring during early plant development gives rise to linked transpositions. In contrast, transposition initiated during reproductive development gives rise to unlinked transpositions in progeny derived from a single F_1_ plant. While linked transpositions initiate when *Ac* transposase is driven from a heterologous promoter such as 35S (**Keller et al., 1993**), unlinked transpositions are derived when *Ac* transposase is expressed using a host plant promoter (**Rommens et al., 1993**). Since we analyzed F_2_ plants from a pooled population of F_1_ plants, a specific distinction about the timing of transposition is not feasible. However, since we obtained both unlinked and linked transposition using *Ac* transposase driven by the 35S promoter, it is likely that the use of the 35S promoter did not hinder unlinked transpositions in tomato.

Theoretically, it is expected that Ds-transpositions should be random across the genome without any bias for specific sites. Apparently, heterochromatin regions of the chromosomes show lower transpositions than the euchromatin region (**Chen et al., 2002; Zhao et al., 2002**). Based on the analyzed population size, the excision and reinsertion frequencies of the *Ds* element were 15% and ∼7.5%, respectively. In this study, several Ds insertions were in the heterochromatic region, discounting any specific bias toward the gene integration site. Similarly, no particular bias was observed for the GC-rich region for the *Ds* transposition, as overall transpositions were interspersed across a wide range of GC content.

Unlike rice or Arabidopsis, tomato genic regions constitute ∼5% of the genome. Consistent with the above, 89.2% of *Ds* insertions were in intergenic regions, and only a few were in the genic region. Many of these were in close proximity to genes with essential functions. Although genic mutations are the primary contributors to mutant phenotypes, emerging evidence indicates that intergenic regions, promoters, UTRs, and intronic mutations also affect phenotypes by influencing gene expression. Therefore, these intergenic insertions can be valuable in altering traits and phenotypes.

With genome editing emerging as a powerful tool to study gene function, it is pertinent to compare it with insertion mutagenesis. Unlike insertion mutagenesis, genome-edited mutagenesis necessitates the transformation of a huge number of gRNA constructs designed a priori to disrupt the function of specific genes in plants. Genome editing primarily results in deletions due to the nature of the editing tools. In maize, most genome-edited genes had deletions (60%) rather than insertions (32.5%) (**Liu et al., 2020**), a feature similar to fast-neutron mutagenesis, which primarily generates insertions and deletions (**Li et al., 2017**). In contrast, *Ds*-induced mutations are mainly knockouts and can be used for activation tagging. Moreover, once a set of starter lines are generated, insertional mutagenesis can be carried out at a large scale without the rigmarole of gRNA design and plant transformation.

In conclusion, we present a two-component *Ac*/*Ds* system for insertional mutagenesis in tomato. Our starter lines are available for distribution to groups interested in generating tomato insertional mutant lines for their studies or for expanding the mutant resources by developing additional lines. By deploying these lines among different groups, a community-wide effort can be made to saturate the tomato genome with insertional mutant lines. A collaborative approach will indubitably enhance the understanding of gene functions and facilitate comprehensive studies of tomato genetics and biology.

## Materials and Methods

### Plant material, plasmids, and growth conditions

Tomato (*Solanum lycopersicum*), cv. Arka Vikas seeds were obtained from the Indian Institute of Horticulture, Bangalore, India. Seeds were surface sterilized with 4% (v/v) sodium hypochlorite for 10 min and then sown in coconut peat (Sri Balaji Agro Services, India). Seedlings were initially grown in a growth room at 25±1°C under 16 h/8 h light and darkness cycles. Afterward, the seedlings were transferred to the greenhouse, and plants were grown under natural day and night cycles. To visualize transgenic seedling fluorescence, the seeds were germinated and grown in a dark room at 25±1°C. The plasmids pSSZ40, pSSZ36, and pSQ3 (**Kolesnik et al., 2004; Qu et al., 2008**) were a gift from Prof. Venkatesan Sundaresan, Department of Plant Biology, University of California, USA.

### Vector construction Preparation of *Ac* vector

The *Ac* vector was made using pBINPLUS as a backbone of the construct and modified from the plasmids pSSZ36 and pSSZ40 **(Figure S2, S3).** Briefly, the CaMV*35S::Ac* cassette was excised from pSSZ36 as a 4.7 Kb BamHI/SalI insert which comprises a minimal *CaMV35S* promoter to drive the expression of the *Ac* gene in plants. The excised *Ac* gene was cloned into the binary vector pBINPLUS (**Figure S3 A-B**). The *ZmUbi1::sGFP* expression cassette was excised as a 3 Kb BamHI/KpnI insert from pSSZ40 and cloned into the backbone of the vector pBINPLUS (**Figure S3 C-G**). The *ZmUbi*1 promoter was replaced with the *CaMV35S* promoter excised from the pBI121 vector using HindIII/XbaI. Promoter replacement was performed to improve reporter gene expression in tomato, a dicot plant. This was followed by cloning a 2 Kb HindIII/KpnI fragment containing a reporter gene cassette (*CaMV35S*::*sGFP*::*NosT*) alongside *Ac* in the backbone of vector pBINPLUS. (**Figure S3 H-K**).

### Preparation of *Ds* vector

The original *Ds* construct was designed for efficient expression in a monocot system. To adapt it for tomato, the *Ds* vector was constructed using pBINPLUS as the backbone and *Ds* from the plasmid pSQ3. A 6.7 kb SacI insert comprising *Ds* 5′ and 3′ sequences bordering 4X *CaMV35S* enhancer:*ZmUbi*1*::RFP/DsRed* was excised from pSQ3 (**Figure S4, S5A-C**) and cloned into the pBINPLUS vector, which already contained the *GFP* cassette (**Figure S5 D-F**). Next, the *ZmUbi1* promoter regulating the expression of *RFP* was replaced with the *AtUbi3* promoter for enhanced expression of the *RFP* gene in tomato.

### Genetic transformation

Tomato genetic transformation was performed using the Agrobacterium*-*mediated cocultivation method with some modifications (**Sharma et al., 2009**). The composition of MS medium and hormones at different stages of regeneration are summarized in **Table S4.** Regenerated plants were acclimatized and grown in pots in the greenhouse until maturity.

### Genomic DNA isolation and PCR analysis

Genomic DNA was isolated from tomato leaves following the protocol described by **Sreelakshmi et al. (2010).** For PCR amplification, 50-100 ng of isolated genomic DNA was used. PCR was carried out using a Bio-Rad thermocycler with specific cycling conditions for each primer set. To confirm the presence of transgenes in the plants, primers specific to transgenes, such as *Ac* and NPTII, were utilized (**Figure S7 A-C**). Additionally, in the T_1_/T_2_/T_3_ generations, a transgene segregation assay was performed using a PCR-based approach, where a forward primer specific to the NPTII gene and reverse primers specific to the *NOS* terminator were used, resulting in the amplification of a ∼750 bp transgene product (**Figure S7**). In the F_2_ generation, *Ac*/*Ds* element segregation was also monitored using GFP- and RFP-specific primers (**Figure 4**). The list of primers used in the study is provided in **Table S5.**

### Southern blot analysis and Multiplex PCR

For Southern blotting, ∼10-12 µg of genomic DNA was digested with the desired restriction enzymes and separated on a 0.8% agarose gel. Southern blotting was then carried out following the method described by **Sambrook et al. (1989).** For transgene screening, a 4.7 Kb BamHI/SalI *Ac* insert (for *Ac* transgenic lines) from pSSZ36 was used as a probe, while for screening the copy number or transgene in transgenic lines (for *Ds* transgenic lines), a 780 bp *NPT*II insert from pBINPLUS was used as a probe (**Figure S8**). Multiplex PCR was performed to rule out the possibility of *Agrobacterium tumefaciens* persistence in transgenic lines (**Figure S9, Table S5),** as described in **Sharada et al. (2017).**

### GFP/RFP fluorescence assay

The *Ac* and *Ds* construct for raising the insertional mutant population have two fluorescent genes, *sGFP/GFP* and *DsRed/RFP,* as visual reporter markers. The putative transgenic plants, untransformed control plants (as a negative control), and tomato homozygous *GFP* plants in the Moneymaker background (**Quadrana et al., 2011**) (as a positive control) were examined for fluorescent protein expression. In the T_0_ generation, leaves of transgenic plants were visualized under a confocal microscope, and images were captured (Confocal Laser Scanning Microscope, Leica TCS SP2). For *sGFP,* excitation/detection at 450/535 nm wavelength was used.

In the T_1_/T_2_/T_3_ generation, 5-day-old dark-grown seedlings of *Ac* and *Ds-*transformed plants were screened for *sGFP* and *DsRed* fluorescence at the Kodak imaging station. For *sGFP,* excitation/emission was at 450/535 nm; for *DsRed,* excitation/emission at 572/610 nm wavelength was used. Similarly, in the F_1_ and F_2_ generations (to detect the presence of only jumped *Ds* elements), seedlings were screened for *sGFP/GFP* and *DsRed/RFP* fluorescence. For fluorescence detection, Brucker MI project software was used. This software gives the image either in black-and-white format or as a colored heatmap.

### Kanamycin Painting

A kanamycin leaf painting assay was performed to check transgene segregation in successive generations and to monitor *in vivo* expression. The leaflets from the same nodes (7^th^ or 8^th^ node) of a one-month-old plant were selected for the kanamycin painting assay from both transgenic lines and wild-type plants. Three leaves of each plant (transgenic and wild type) were tagged, and the upper surface of the leaves was painted with 250 µg/mL (w/v) kanamycin solution. After ten days of incubation, leaves were observed for the appearance of the bleached phenotype.

### Determination of the T-DNA insertion sites

For retrieval of flanking sequences, we used a modified version of TAIL-PCR, i.e., FPNI-PCR (Fusion primer and nested integrated PCR) (**Wang et al., 2011**). Sequences flanking *Ds* insertion in the genome were amplified using primers and thermocycler protocols with some modifications as published in **Sharada et al. (2017)** (**Table S6**). Filtering alignment and analysis of the sequenced product were performed as described in **Sharada et al. (2017).** For retrieval of *Ds*-transposition sites, inverse PCR using the protocol of **Han et al. (2018)** was also used (**Table S7**).

## Declarations

### Ethics approval and consent to participate

Not applicable

### Consent for publication

All authors have approved the manuscript and given consent for its publication.

### Availability of data and material

The *Ac*/*Ds* starter lines are available for distribution on executing the Material Transfer Agreement with the Repository of Tomato Genomics Resources, University of Hyderabad, India.

### Author Contributions

YS and RS designed this project and wrote the manuscript. AK, RP, and AP together performed the experiments and data analysis. All authors read and approved the manuscript.

## Acknowledgments

We thank Prof. Venkatesan Sundaresan, Department of Plant Biology, University of California, USA. For providing is the plasmids pSSZ40, pSSZ36, and pSQ3. We thank Dr. Imran Siddiqui, Center of Cellular and Molecular Biology, Hyderabad, India, for helpful discussions and support. We thank Dr. Sanchri Sircar for her assistance in the analysis of gene insertion sites.

---

**Figure S1.**
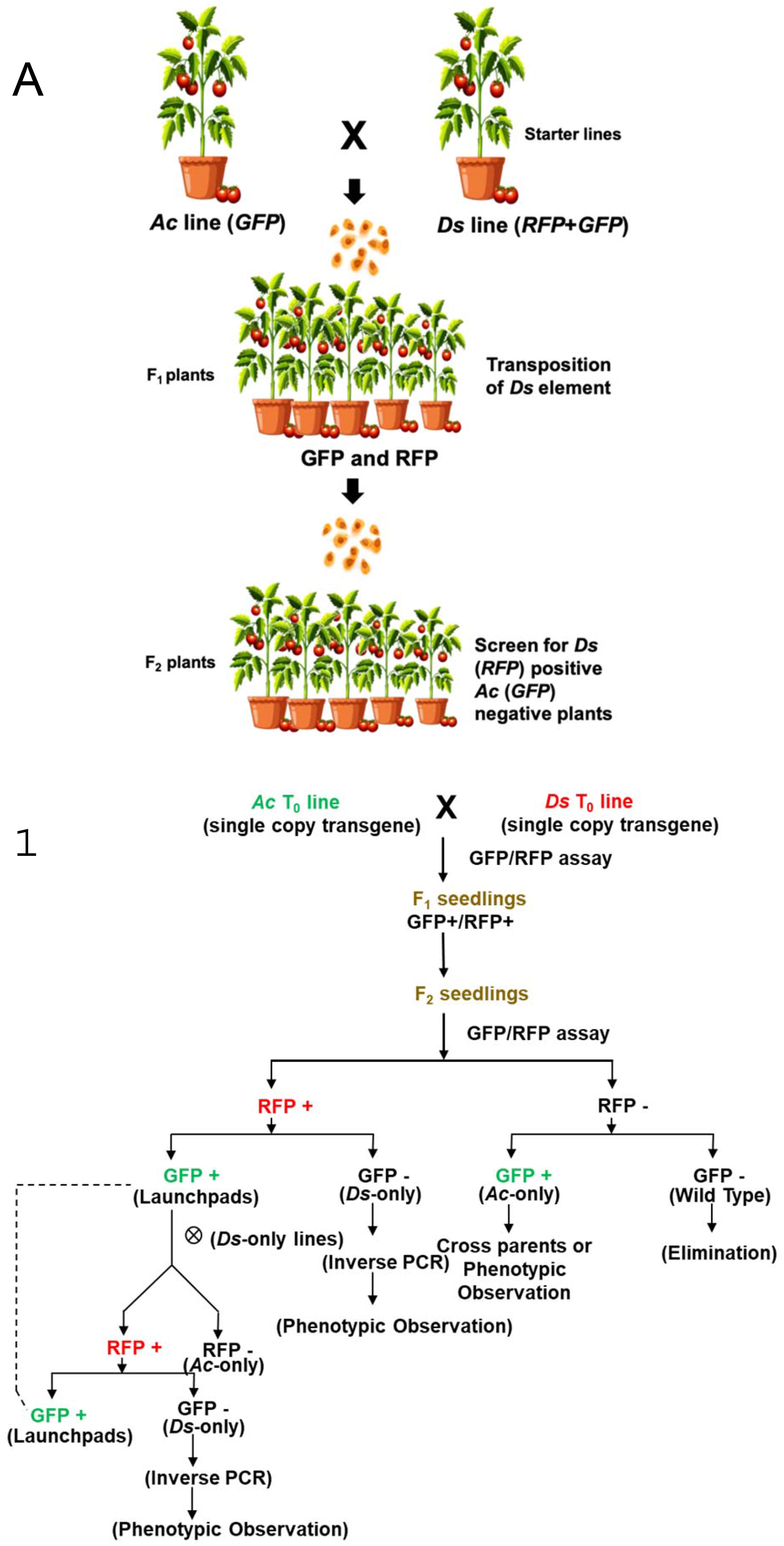
The screening process for selecting plants with *Ds* transposition. **A.** Graphical illustration of generation of transposon-tagged mutant population in tomato using *Ac/Ds* transposon system. **B.** The flowchart outlines the screening process for selecting plants with *Ds* transposition. The presence of both GFP and RFP genes in successive generations is assessed through fluorescence assay and PCR analysis. The F_1_ seedlings carrying both *Ac* and *Ds* elements (GFP+/RFP+) are carried forward to the F_2_ generation. Within the F_2_ population, based on screening plants are classified into four categories. Plants exhibiting only RFP (GFP-/RFP+) indicate successful transposition of *Ds* into a new site. Plants displaying only GFP (GFP+/RFP-) are identified as *Ac*-only plants. Wild-type plants are characterized by neither GFP nor RFP (GFP-/RFP-). Plants exhibiting both GFP and RFP (GFP+/RFP+) can serve as “launch pads” for identifying new instances of *Ds* transposition, as described above.

**Figure S2.**
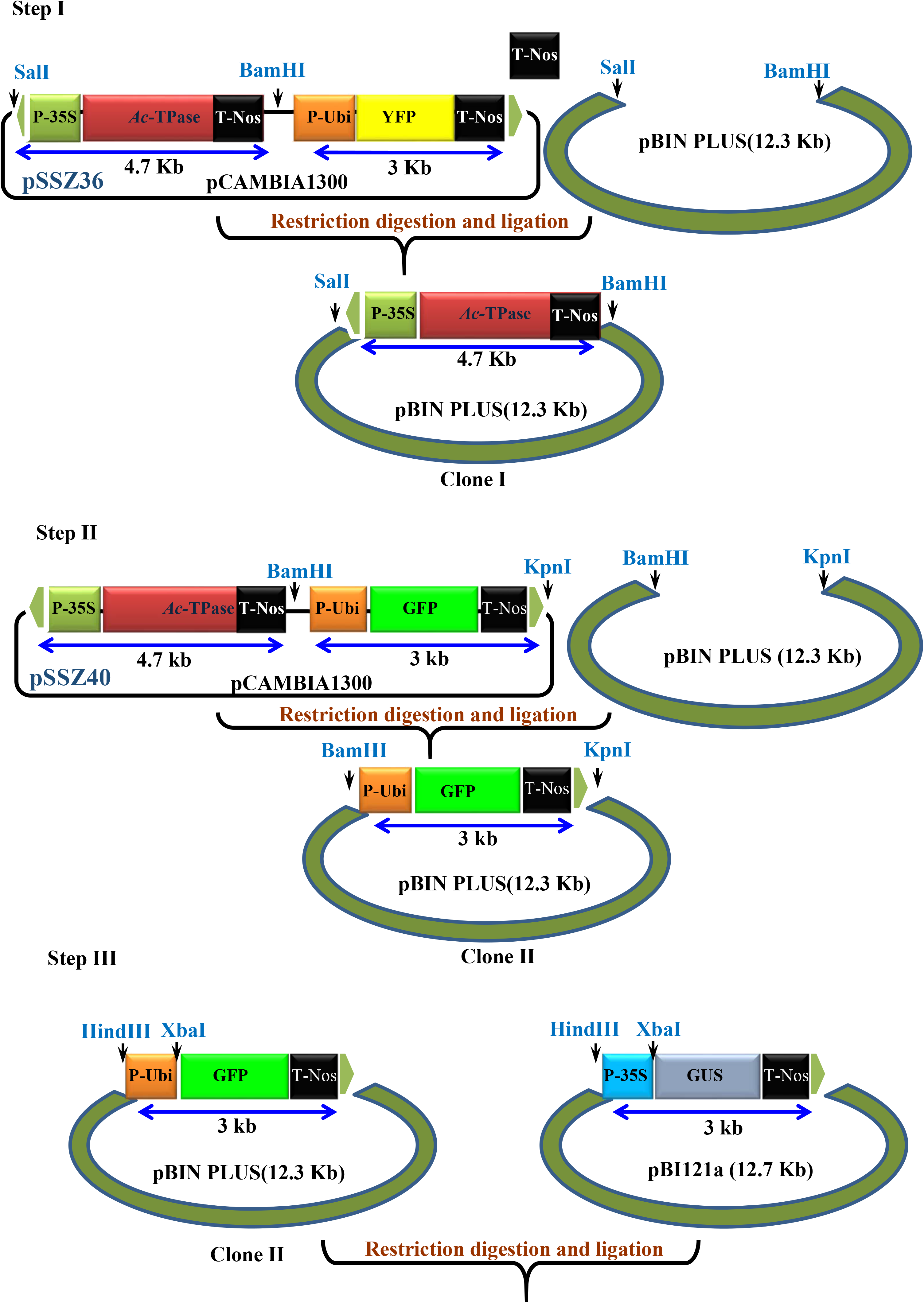
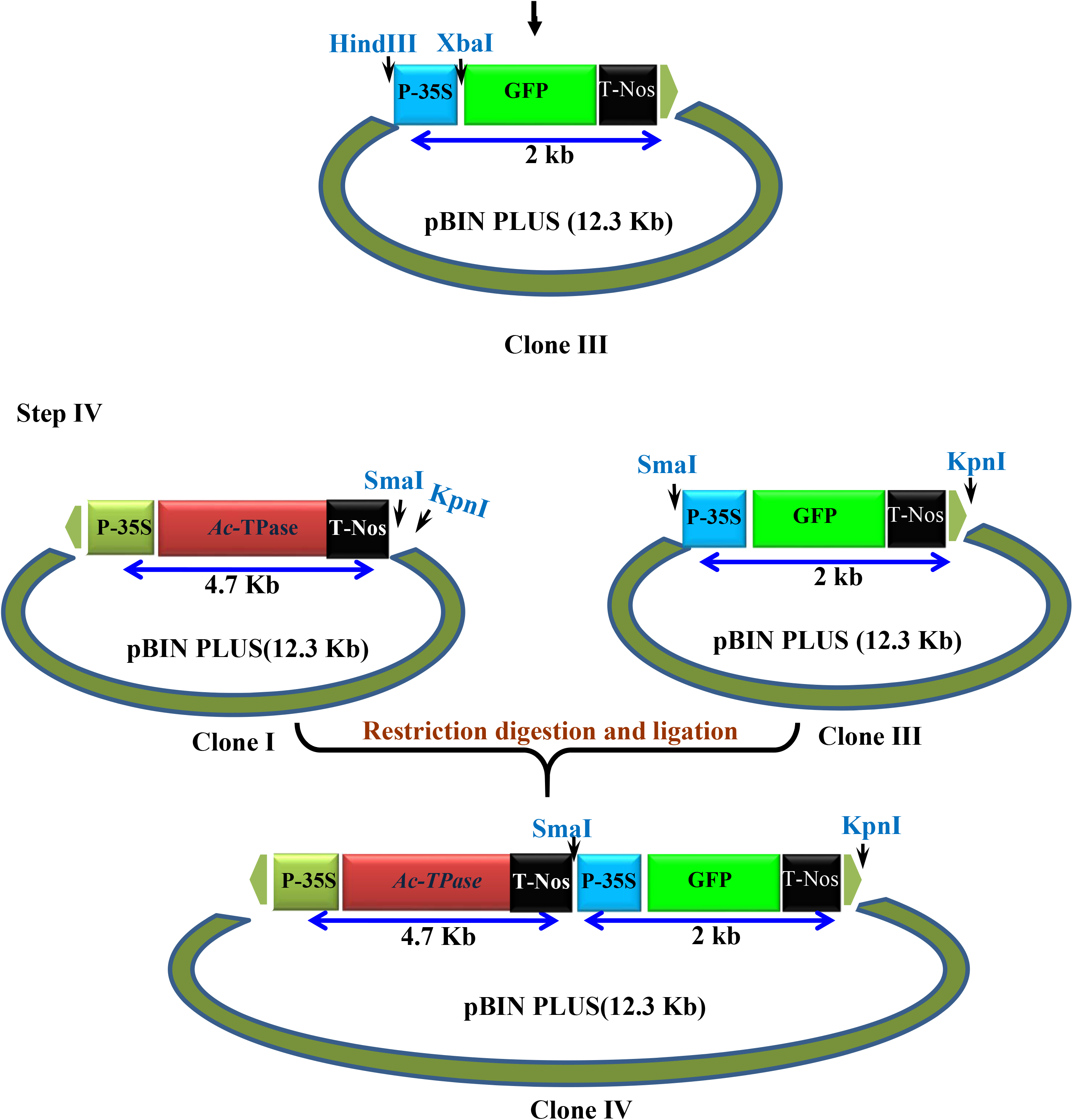
A schematic representation of the steps involved in mobilizing the *Ac-Tpase (pSSZ36)* construct into the *pBINPLUS* plasmid. The scheme also illustrates the replacement of the maize ubiquitin promoter with the *35S* promoter to drive *GFP* expression (*pSSZ40*), followed by the final mobilization of the modified construct into the *pBINPLUS* plasmid alongside the *Ac-TPase* construct.

**Figure S3.**
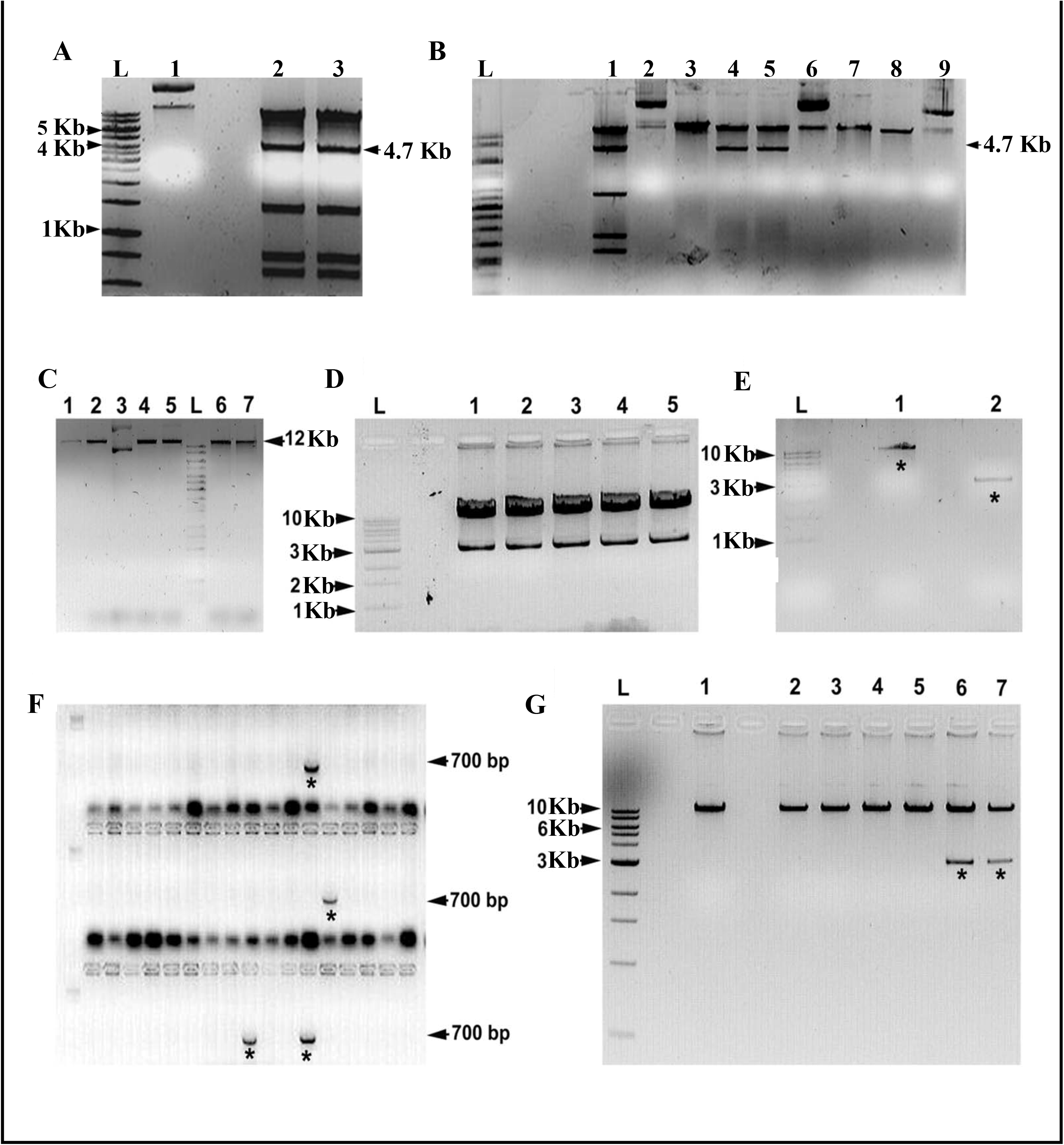
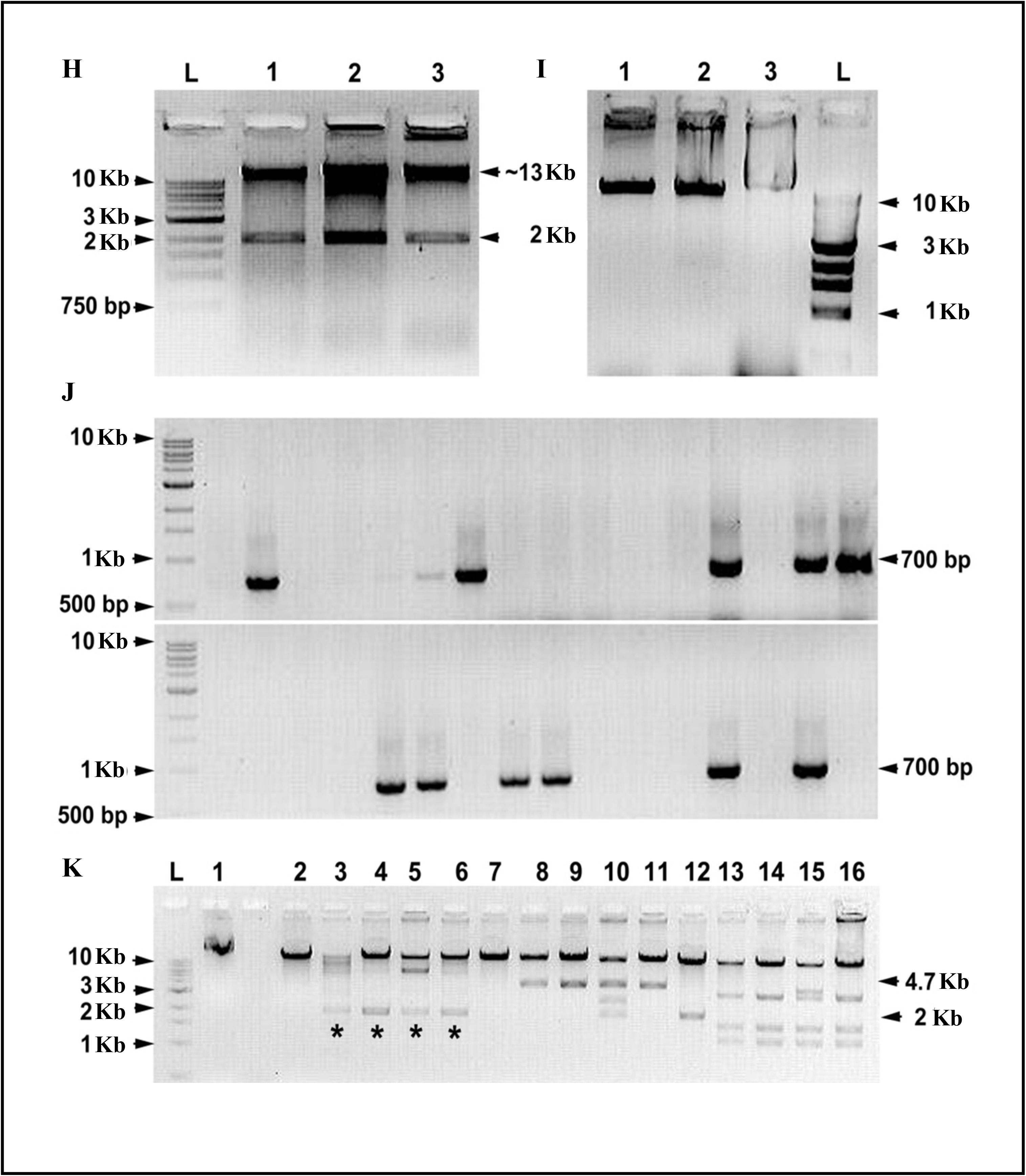
The generation of the Ac-TPase constructs through various molecular steps. **A.** Restriction digestion of *pSSZ36* vector. Lane 1, Undigested plasmid; Lane 2-3, Plasmid digested with BamHI/SalI. **B.** Restriction digestion of recombinant *pBINPLUS* plasmid and <u>pSSZ36</u> bearing plasmid. Lane 1, *pSSZ36* plasmid digested with BamHI/SalI showing the release of 4.7 Kb band and three additional bands; Lane 2, Undigested *pSSZ36* plasmid; Lane 3, Recombinant *pBINPLUS* plasmid digested with BamHI; Lane 4-5, Release of 4.7 Kb fragment from recombinant *pBINPLUS* plasmid digested with BamHI/SalI (**Note**: Only one 4.7 Kb band is released in compared to *pSSZ36* that shows four bands along with 4.7 kb band); Lane 6, Undigested recombinant *pBINPLUS* plasmid; Lane 7-8, *pBINPLUS* digested with BamHI/SalI; Lane 9, Undigested *pBINPLUS* plasmid. **C.** Restriction digestion profile of *pBINPLUS*; Lane 1-2, Plasmid digested with BamHI; Lane 3, Undigested plasmid; Lane 4-5, Plasmid digested with KpnI; Lane 6, Plasmid digested sequentially with BamHI followed by KpnI; Lane 7, Plasmid digested sequentially with KpnI followed by BamHI. **D.** Lane 1-5, *pSSZ40* vector digested with BamHI/KpnI. **E.** Gel eluted plasmids digested with BamHI/KpnI; Lane 1, Linearized *pBINPLUS* plasmid; Lane 2, 3 Kb cassette from pSSZ40 plasmid. **F.** Colony PCR using *GFP-*specific primers; **Note,** Few colonies showed the presence of a 700 bp product. **G.** Restriction digestion of plasmid (*pBINPLUS* having *ZmUbi+sGFP*+*NOSter* cassette) isolated from PCR positive colonies; Lane 1-7, transformed *pBINPLUS* plasmid digested with BamHI/KpnI. Lanes 6-7 show the release of a 3 Kb *GFP* reporter cassette. **\***Mark in each gel shows the amplified PCR product or release of the desired insert. **H.** Release of *GFP* reporter gene cassette from *pBINPLUS* upon digestion with SmaI and KpnI; Lane 1-3 show release of 2 Kb reporter gene cassette. **I.** Restriction profile of *pBINPLUS* having *Ac-TPase* element. Lane 1-2, Linearized plasmid after digestion with SmaI and KpnI; Lane 3, Undigested plasmid. (**Note**: Linearized plasmid in Lane 1-2 has more gel mobility than undigested plasmid). **J.** Colony PCR using *GFP-specific* primers. The presence of insert was identified by PCR amplification of a 700 bp product. **Note**: Few colonies show the presence of a 700 bp product. **K.** Restriction profile of cloned (*sGFP* reporter gene cassette with *Ac-TPase* element in *pBINPLUS* plasmid); Lane 1, Undigested plasmid; Lane 2, Linearized *pBINPLUS* with *Ac-TPase* element after digestion with SmaI and KpnI; Lane 3-6, Release of 2 Kb *GFP* reporter cassette after digestion with SmaI and KpnI (marked with*); Lane 7, Linearized *pBINPLUS* with *GFP* reporter cassette after digestion with BamHI/SalI; Lane 8-11, Release of 4.7 Kb *Ac-TPase* after digestion with BamHI/SalI; Lane 12, Digestion of *pBINPLUS* plasmid with *GFP* reporter cassette with HindIII; Lane 13-16, Recombinant clones digested with HindIII enzymes **Note**: *Ac-TPase* has five internal restriction sites for HindIII enzymes. In each gel, **Lane L** 1 Kb DNA ladder.

**Figure S4.**
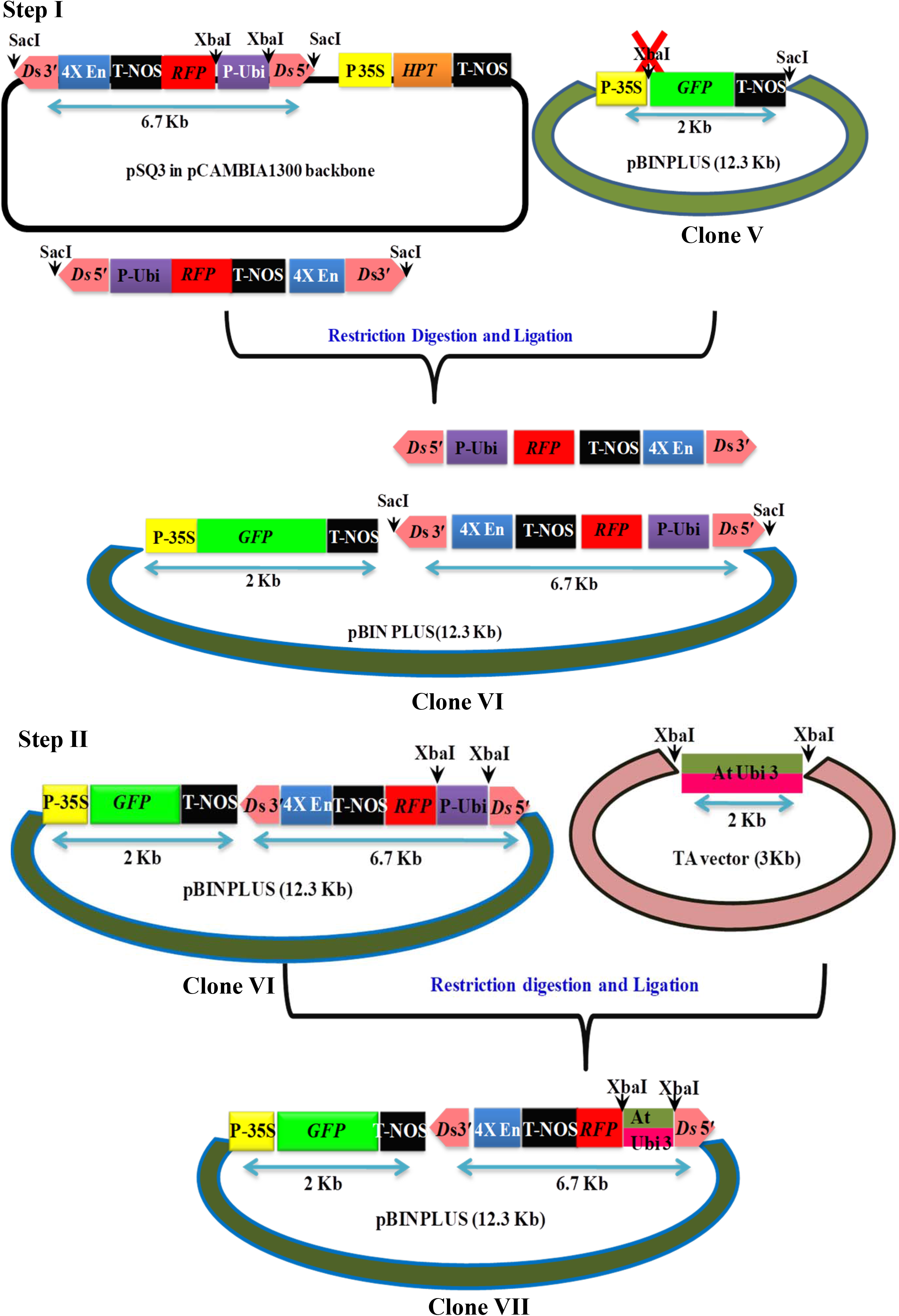
A schematic representation of the steps involved in mobilizing the *Ds (pSQ3)* construct into the *pBINPLUS* plasmid. The scheme illustrates the replacement of the maize ubiquitin promoter with the Arabidopsis ubiquitin promoter to drive *Ds* expression, and replacement of the hygromycin gene with GFP, and the mobilization of the modified construct into the *pBINPLUS* plasmid.

**Figure S5.**
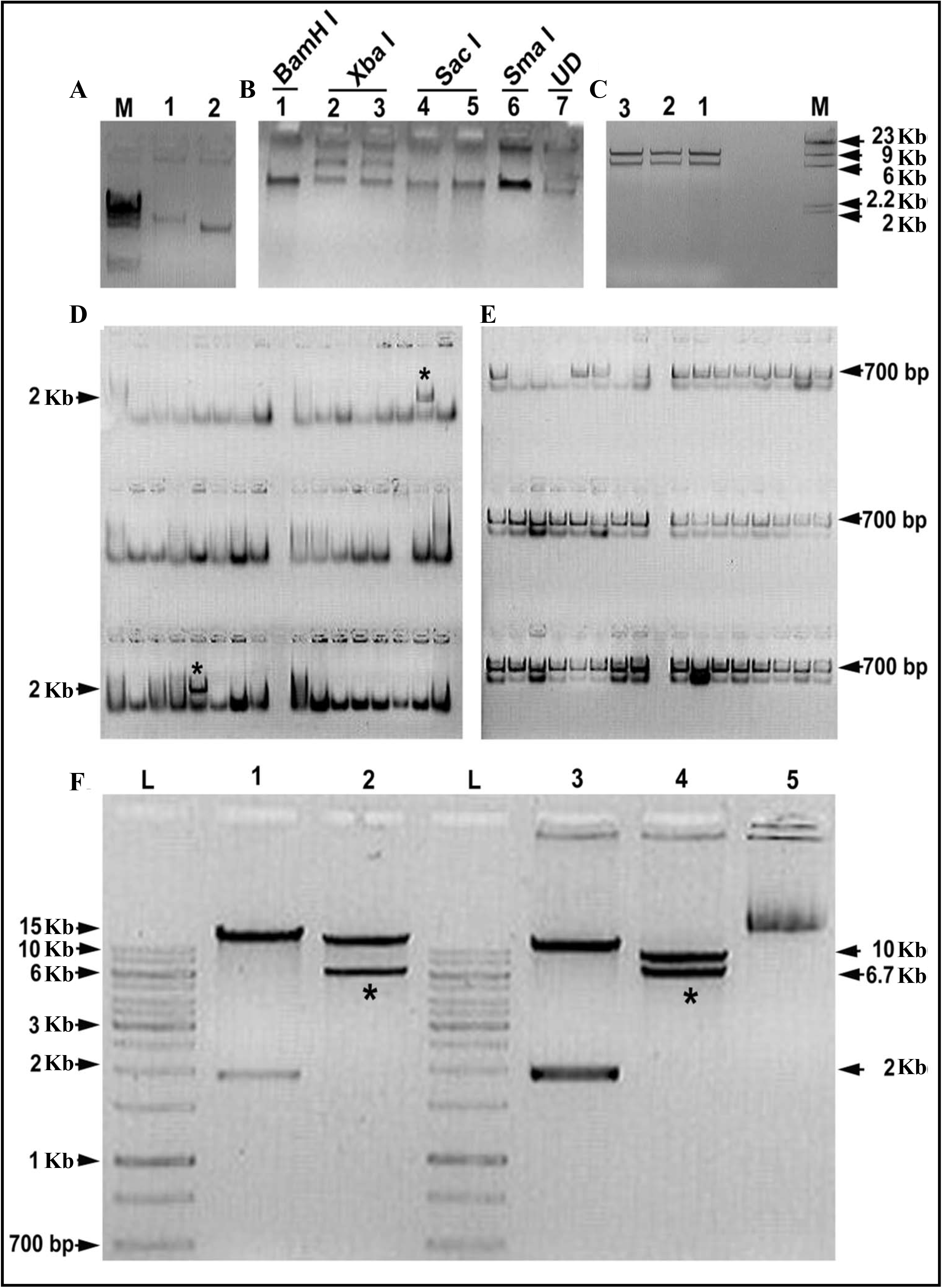
Generation of *Ds* construct. **A.** Linearization of *pBINPLUS* plasmid harboring *GFP* reporter cassette with XbaI. Lane 1, Plasmid digested with XbaI; Lane 2, Undigested plasmid; Lane M, λ DNA/ HindIII digested ladder. **B.** Confirmation of disruption of XbaI restriction site in *pBINPLUS* plasmid harboring *GFP* reporter cassette. Lane 1, Linearized plasmid after digestion with BamHI enzyme, Lane 2-3, Undigested plasmid after digestion with XbaI enzyme; Lane 4-5, Linearized plasmid after digestion with SacI enzyme; Lane 6, Linearized plasmid after digestion with SmaI enzyme; Lane 7, Undigested plasmid. **C.** Release of *Ds* cassette from pSQ3 plasmid; Lane 1-3, 6.7 Kb fragment released from pSQ3 after digestion with SacI enzyme; Lane M, λ DNA/ HindIII digested ladder. **D.** Colony PCR of recombinant colonies with *ZmUbi* promoter-specific primers, **Note:** Amplicon of a 2 Kb size (*) indicates the presence of insert. **E.** Colony PCR with *GFP*-specific primers, **Note:** Amplicon of 700 bp size (*) indicates the presence of backbone. **F.** Mobilization of 6.7 Kb *Ds* element cassette to pBINPLUS binary vector with *GFP* reporter gene. Lane 1, Release of 2 Kb *ZmUbi* promoter from pBINPLUS plasmid after digestion with XbaI; Lane 2, Release of 6.7 Kb *Ds* cassette from pBINPLUS after digestion with SacI; Lane 3, Release of 2 Kb *ZmUbi* promoter from pSQ3 plasmid after digestion with XbaI; Lane 4, pSQ3 plasmid digested with SacI releases a 6.7 Kb fragment along with 10 Kb backbone; Lane 5, Undigested pSQ3 plasmid; **Lane L,** 1 Kb DNA ladder in each gel.

**Figure S6:**
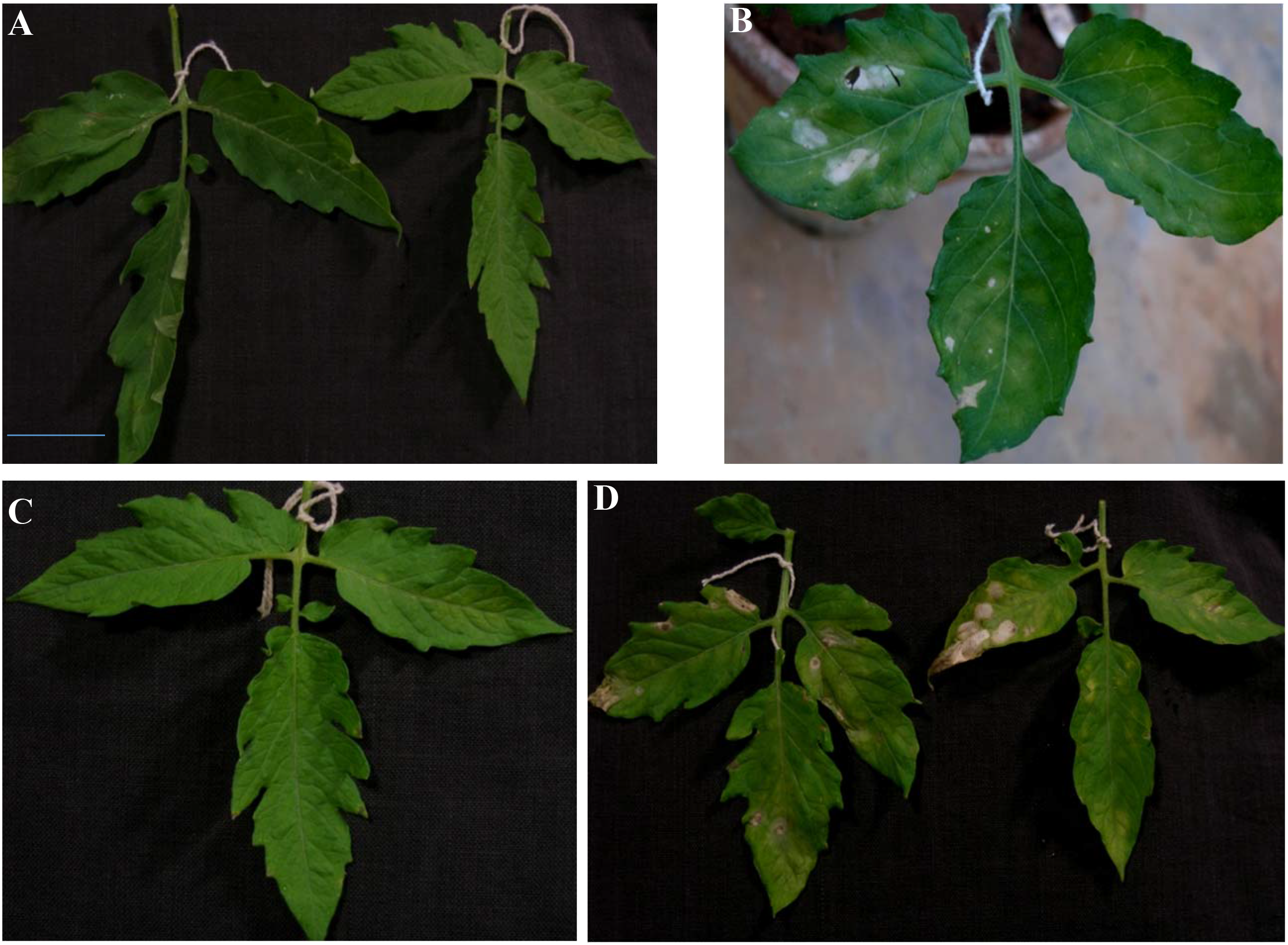
Kanamycin painting assay for identification of transgenic plants. **A-B.** The wild-type leaf after ten days of mock painting (**A**) and kanamycin painting (**B**). **C.** Leaf of a transgenic line exhibiting resistance to kanamycin. **D.** Leaf of a transgenic line exhibiting sensitivity to kanamycin. Note the appearance of bleached spots at the point of kanamycin application.

**Figure S7:**
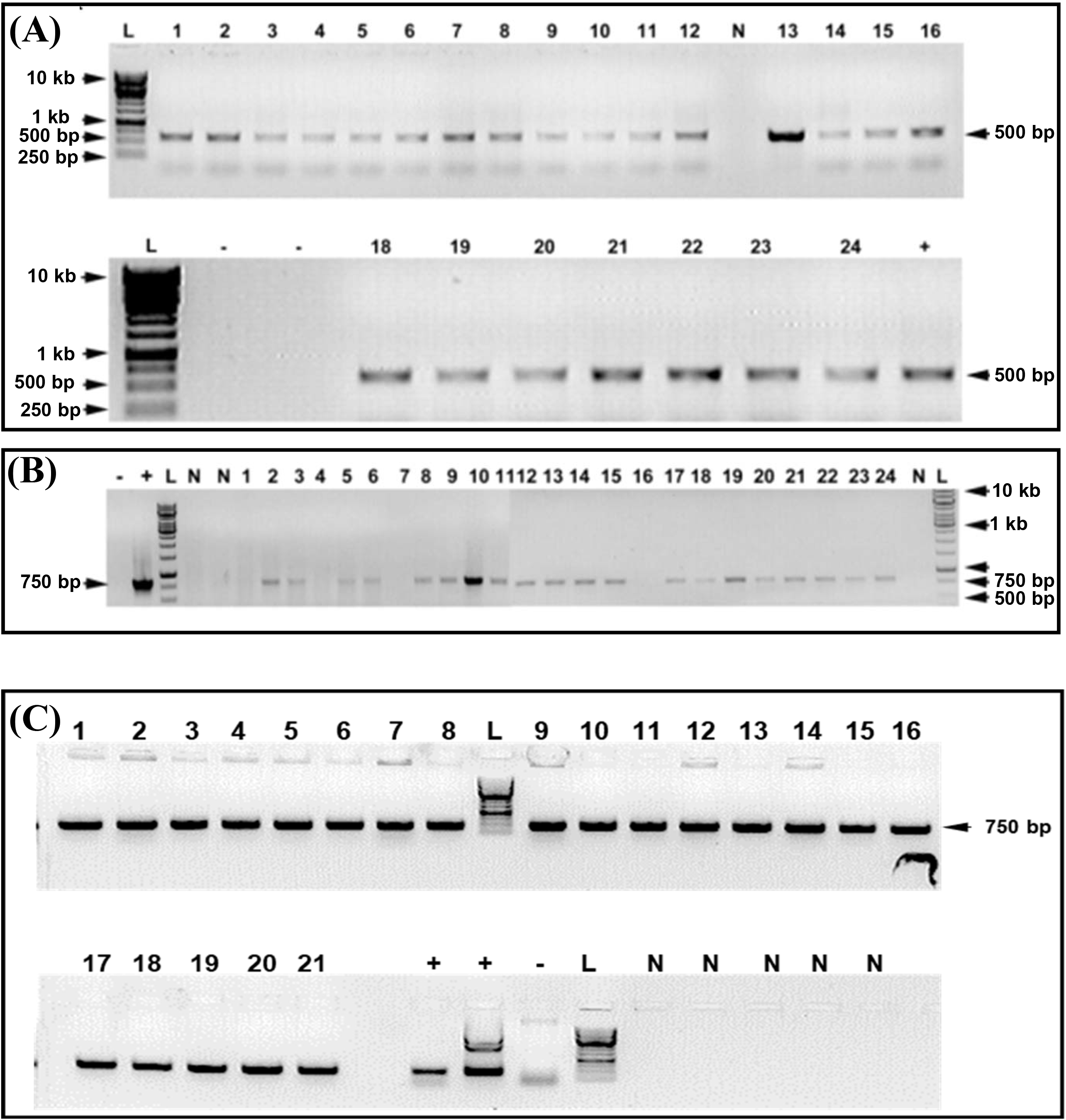
Detection of the transgene in *Ac*-*TPase* and *Ds* T_0_ plants by PCR. **A**. PCR with primers specific to *Ac*-*TPase* shows amplification of ∼500 bp product of transgene. **B-C**. PCR with primers specific to *NPTII* shows amplification of ∼750 bp product in *Ac*-*TPase* and *Ds* plants, respectively. In each gel, Lane L signifies a 1 Kb DNA ladder. Symbols **+** and **–** designate positive (**+**) and negative control (**-**). Plasmid DNA of the construct employed in the transformation was used as positive control and the genomic DNA of the untransformed plant was used as the negative control. **N** designates as an empty lane. Numbers on each lane indicate the line number of the respective T_0_ plant.

**Figure S8:**
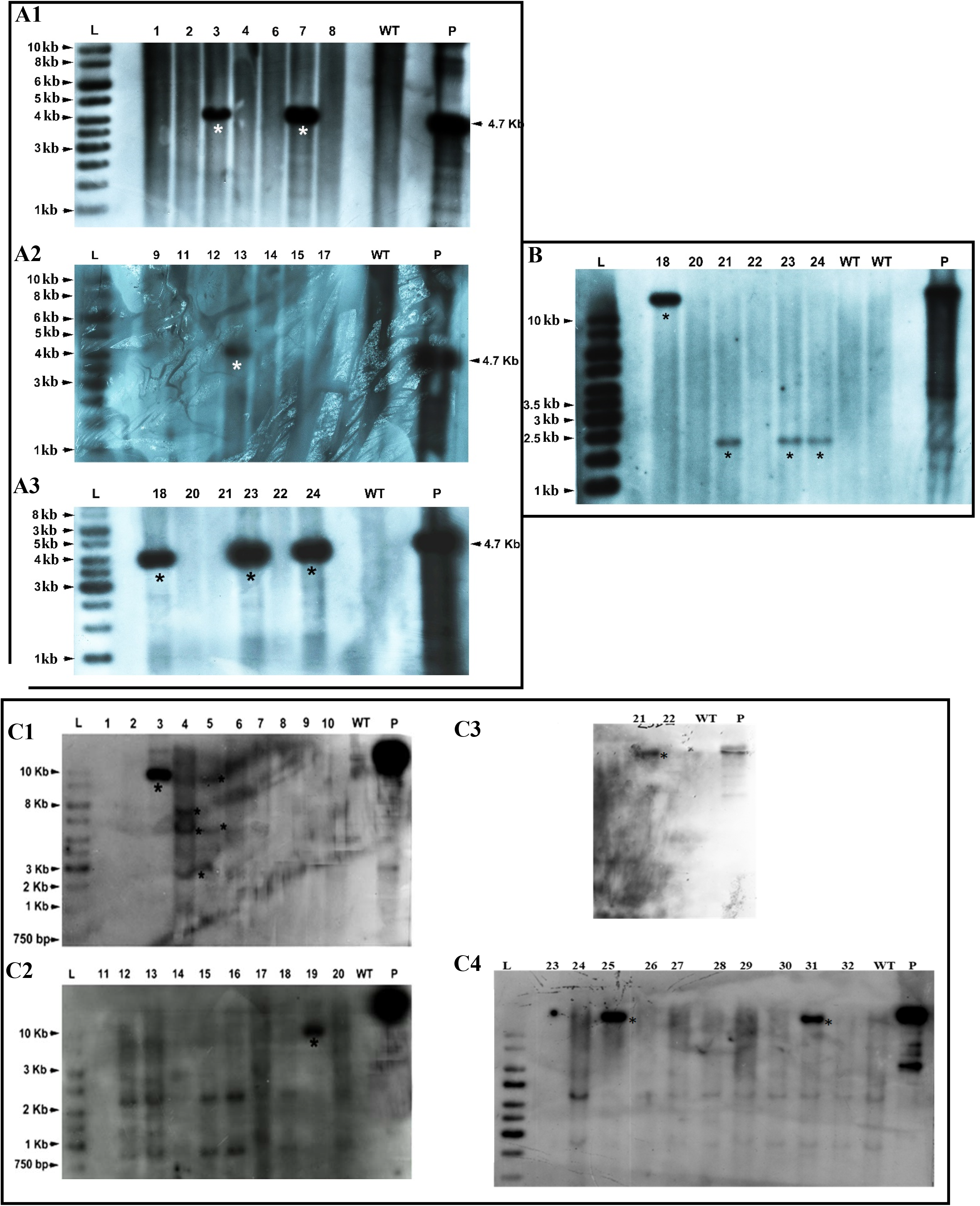
Southern blots of *Ac-TPase* lines for ascertaining the transgene presence and copy number. **A1-A3**. For monitoring the transgene integration in transformed lines, a 4.7 kb (*Ac-TPase* gene) fragment was used as a probe. The genomic DNA of T_0_ plants was digested with BamHI/SalI. **B.** For checking the copy number of the transgene. The genomic DNA of *Ac-TPase* T_0_ plants was digested with BamHI and probed with the radiolabelled *NPTII* gene. **C1-C4.** Genomic DNA of *Ac-TPase* T_2_ plants digested with BamHI. The blot was probed with a radiolabelled *NPTII* gene. Lane **L**, 1 kb DNA Ladder; P, Plasmid DNA of the construct used for transformation; WT, Wild type genomic DNA (negative control). **Note:** Lanes showing the presence of transgene in the Transgenic lines are marked with a star (*) symbol. The numbers on top of each lane indicate the line number of the respective T_0_ /T_2_ plant. **Note:** Star mark (*) indicates the presence of *Ac-TPase* transgene in transgenic lines.

**Figure S9:**
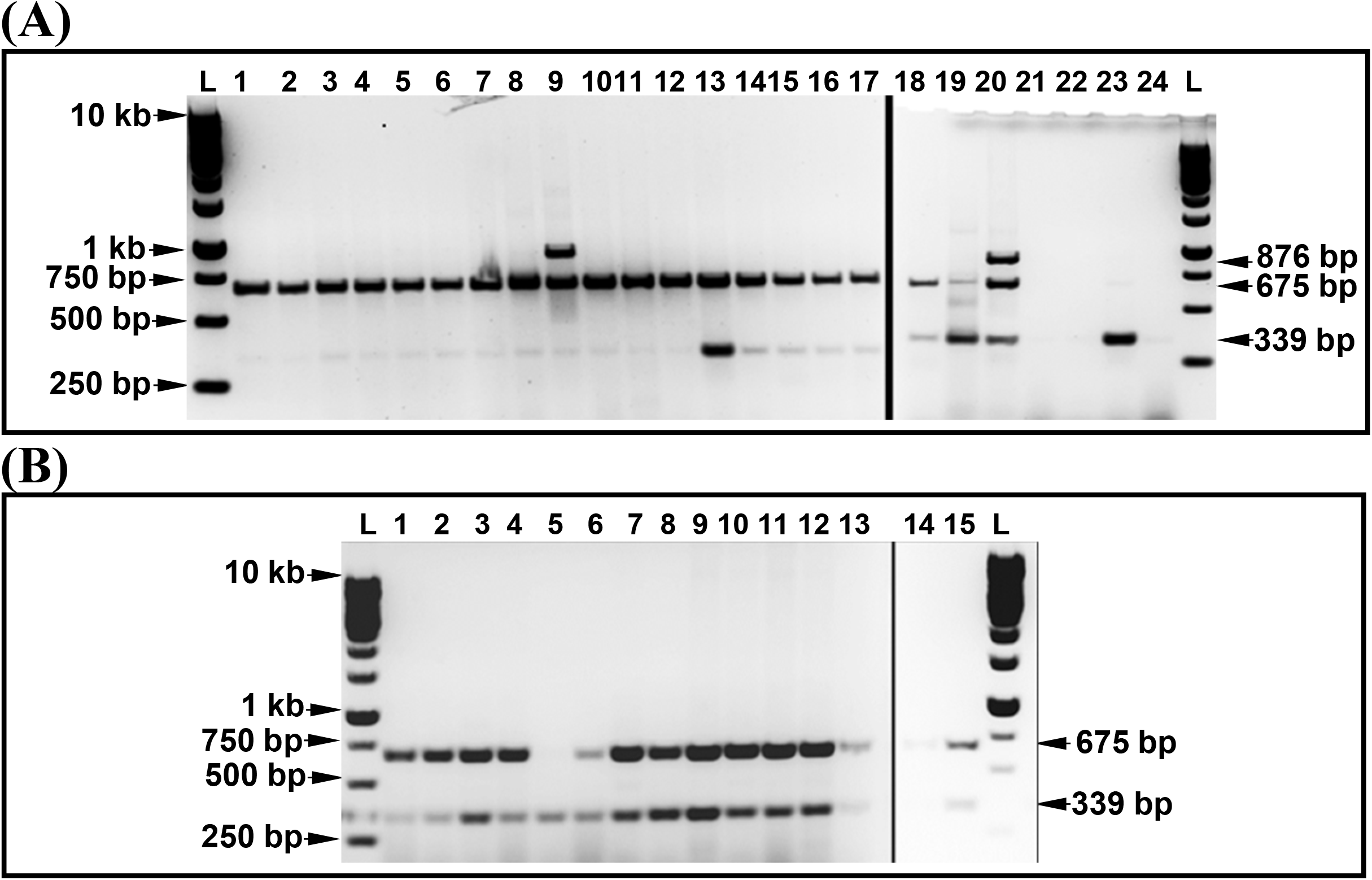
Multiplex PCR to detect the *Agrobacterium* contamination and transgene in *Ac-TPase* and *Ds* tagged Parental lines (T_0_). **A.** Lane **L**,1 kb ladder; Lane 1- 19 *Ds* T_0_ Plants; Lane 20, *Agrobacterium* transformed with *Ds* plasmid; Lane 21, Untransformed *Agrobacterium* strain C58C1; Lane 22, Arka Vikas (WT); Lane 23, Plasmid DNA of *Ds* construct; Lane 24, NTC; Genomic DNA of the untransformed plant (Arka Vikas-AV) was used as a negative control. **B.** Lane L,1 kb ladder; Lane 1- 13, *Ac-TPase* T_0_ Plants; Lane 14, AV (WT); Lane 15, Plasmid DNA of *Ds* construct. **Note**: *chv* gene (876 bp) is *Agrobacterium* specific, whereas *cyc*B (675 bp) is a single copy native gene of tomato, and *NPT*II (339 bp) is a transgene. Numbers on each lane indicate the line number of respective T_0_ plants.

**Figure S10.**
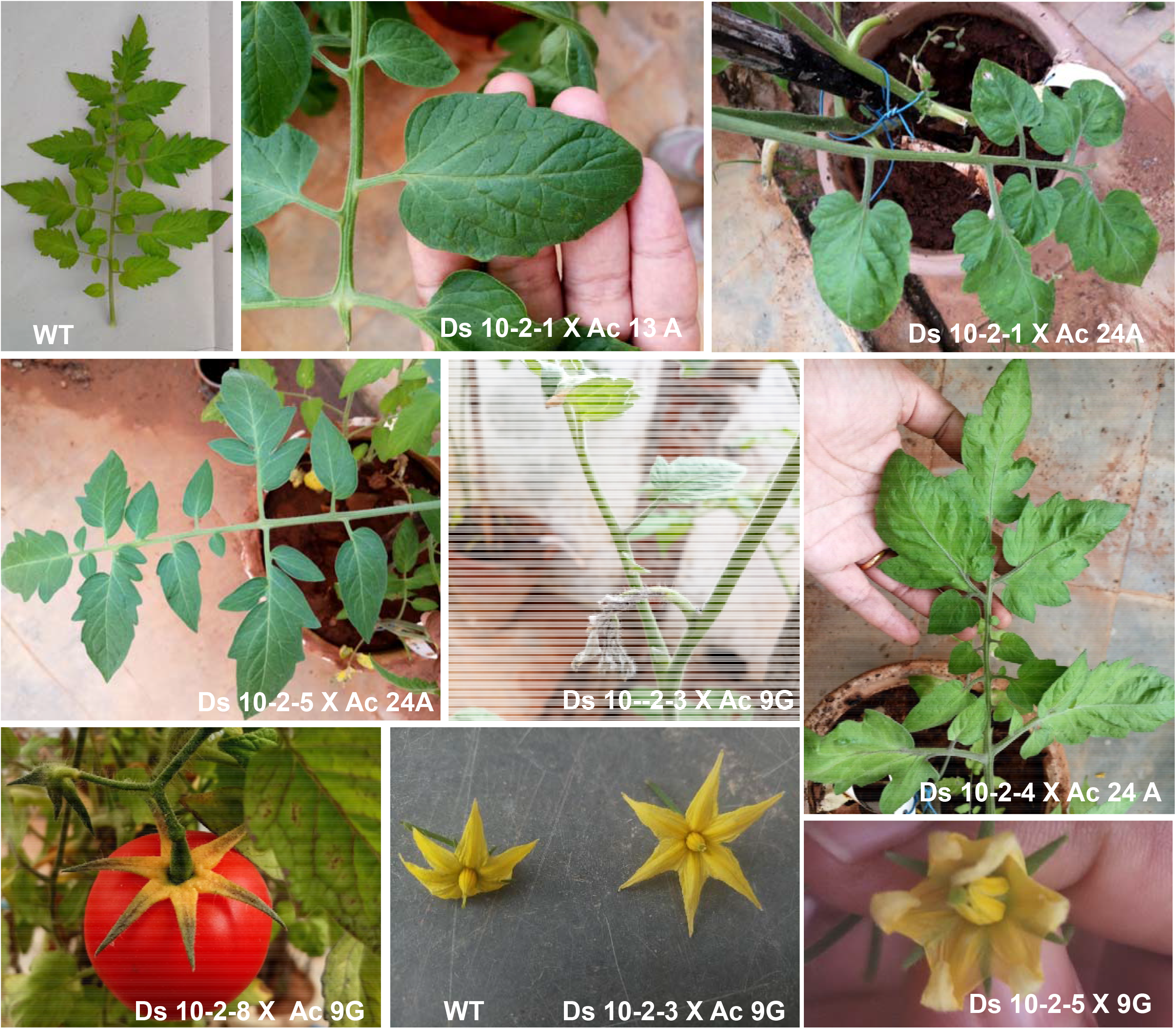
Phenotypic alterations in F_2_ progeny of *Ac* X *Ds* crosses compared to Arka Vikas (WT). The number on the bottom of each picture indicates the name of the cross between *Ac* and *Ds* parents **WT**, Untransformed plant. Note potato leaf phenotypes, leathery leaf phenotypes, fleshy sepals, introse stamen, and altered anther cone structure.

**Figure S11.**
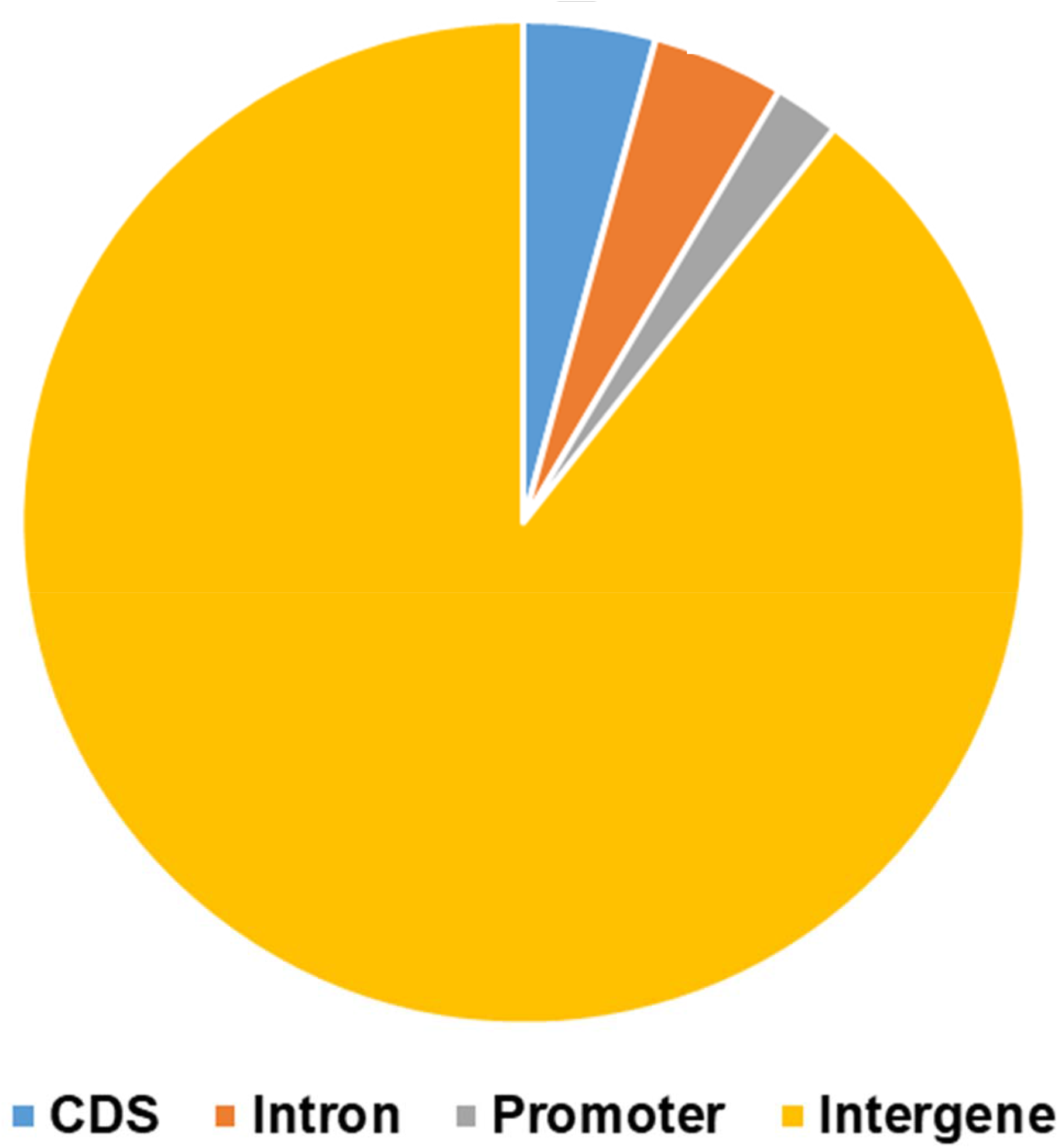
Distribution of *Ds*-insertion sites in Intergenic and Intragenic regions.

**Table S1:** Kanamycin resistance assay of *Ac-TPase* starter lines.

| Line Number | Kanamycin Sensitive | Kanamycin Resistant | PCR Positive lines | Southern Positive lines |
| --- | --- | --- | --- | --- |
| *Ac-TPase* - 1A | *Ac-TPase* - 1A1, 1A2, 1A3, 1A4, 1A7 | *Ac-TPase* - 1A5,1A6 | *Ac-TPase* - 1A5, 1A6 | *Ac-TPase* - 1A6 |
| *Ac-TPase* - 2C | *Ac-TPase* - 2C3 | *Ac-TPase* - 2C1, 2C2, 2C4, 2C5, 2C6, 2C7, 2C8, 2C9, 2C10 | *Ac-TPase* - 2C1, 2C2, 2C6, 2C8, 2C9, 2C10 | *Ac-TPase* - 2C1, 2C2 |
| *Ac-TPase* - 3A | *Ac-TPase* - 3A1, 3A4 | *Ac-TPase* - 3A2, 3A3, 3A5, 3A6 | *Ac-TPase* - 3A2,3A3,3A6 |  |
| *Ac-TPase* - 7B | *Ac-TPase* - 7B1, 7B5, 7B6 | *Ac-TPase* - 7B3, 7B7, 7B9 | *Ac-TPase* - 7B3, 7B7,7B9 |  |
| *Ac-TPase* - 9G | *Ac-TPase* - 9G3, 9G5, 9G6, 9G8 | *Ac-TPase* - 9G1, 9G2, 9G4, 9G7, 9G9, 9G10 | *Ac-TPase* - 9G1, 9G2, 9G9,9G10 | *Ac-TPase* 9G9 |
| *Ac-TPase* - 13A | *Ac-TPase* - 13A7 | *Ac-TPase* - 13A1, 13A2, 13A3, 13A4, 13A5, 13A6 | *Ac-TPase* - 13A1, 13A2, 13A3, 13A5 |  |
| *Ac-TPase* - 13D | *Ac-TPase* - 13D3, 13D4 | *Ac-TPase* - 13D1, 13D2, 13D5, 13D6, 13D7, 13D8 | *Ac-TPase* - 13D1, 13D2, 13D5, 13D7 |  |
| *Ac-TPase* - 21A | *Ac-TPase* - 21A1, 21A2, 21A4 | *Ac-TPase* - 21A3, 21A5, 21A6 | *Ac-TPase* - 21A3, 21A5, 21A6 |  |
| *Ac-TPase* - 23D | *Ac-TPase* - 23D3 | *Ac-TPase* - 23D1, 23D2, 23D4, 23D6, 23D7, 23D9 | *Ac-TPase* - 23D1, 23D2, 23D6, 23D7, 23D9 | *Ac-TPase* - 23D2, 23D6 |
| *Ac-TPase* - 24A | *Ac-TPase* - 24A1, 24A2 | *Ac-TPase* - 24A3,24A4 | *Ac-TPase* - 24A3, 24A4 | *Ac-TPase* - 24A3 |

**Table S2:** The chromosomal location of the *Ds* launching pad.

| S.No. | Name of *Ds* launch pad | Chromosome number | Location in the genome | Location/Sequence ID |
| --- | --- | --- | --- | --- |
| 1. | *Ds-2* | 1 | Zinc finger protein VAR3 gene | Solyc01g099230 |
| 2. | *Ds-4* | 9 | Tic 22-like gene | Solyc09g092530 |
| 3. | *Ds-10-2* | 3 | Intergenic | SL3.0ch03, 26676740-26676787 |

#
#

**Table S3.** Distribution of Ds-tagged lines on different chromosomes of tomato.

| Chromosome number | Number of Ds transpositions |
| --- | --- |
| 1 | 0 |
| 2 | 4 |
| 3 | 21 |
| 4 | 2 |
| 5 | 1 |
| 6 | 2 |
| 7 | 4 |
| 8 | 9 |
| 9 | 3 |
| 10 | 7 |
| 11 | 2 |
| 12 | 1 |
| **Total** | **56** |

#
#

**Table S4:** Composition of MS media used in various transformations and regeneration steps.

| **Components** | **Seed germination** | **Pre-culture and co-cultivation** | **Selection** | **Rooting** |
| --- | --- | --- | --- | --- |
| MS Salts | 0.5 X | 1 X | 1 X | 1 X |
| B5 Vitamins | 0.5 X | 1 X | 1 X | 1 X |
| Sucrose (g/L) | 15 | 30 | 30 | 30 |
| Agar (% w/v) | 0.8 | 0.8 | 0.8 | 0.8 |
| BAP mg/L | 0 | 2 | 0 | 0 |
| IAA mg/L | 0 | 0 | 0.1 | 0 |
| Zeatin mg/L | 0 | 0 | 1 | 0 |
| Kanamycin mg/L | 0 | 0 | 100 | 100 |
| Cefotaxime mg/L | 0 | 0 | 500 | 500 |

**Table S5:**
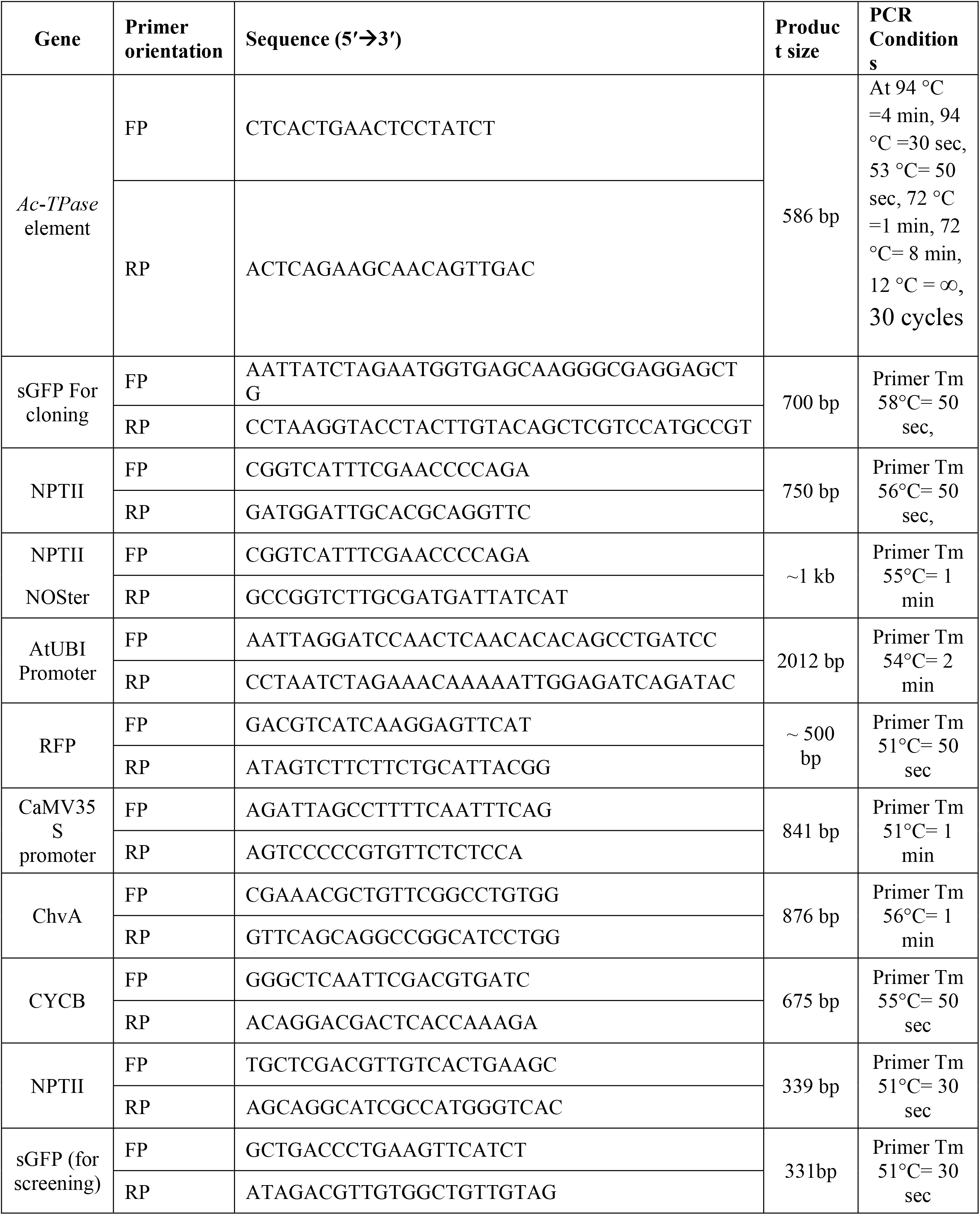
Sequences of primers used for PCR. The PCR product size of the respective primer pair and cycling conditions are also mentioned next to the respective primer pairs.

**Table S6:**
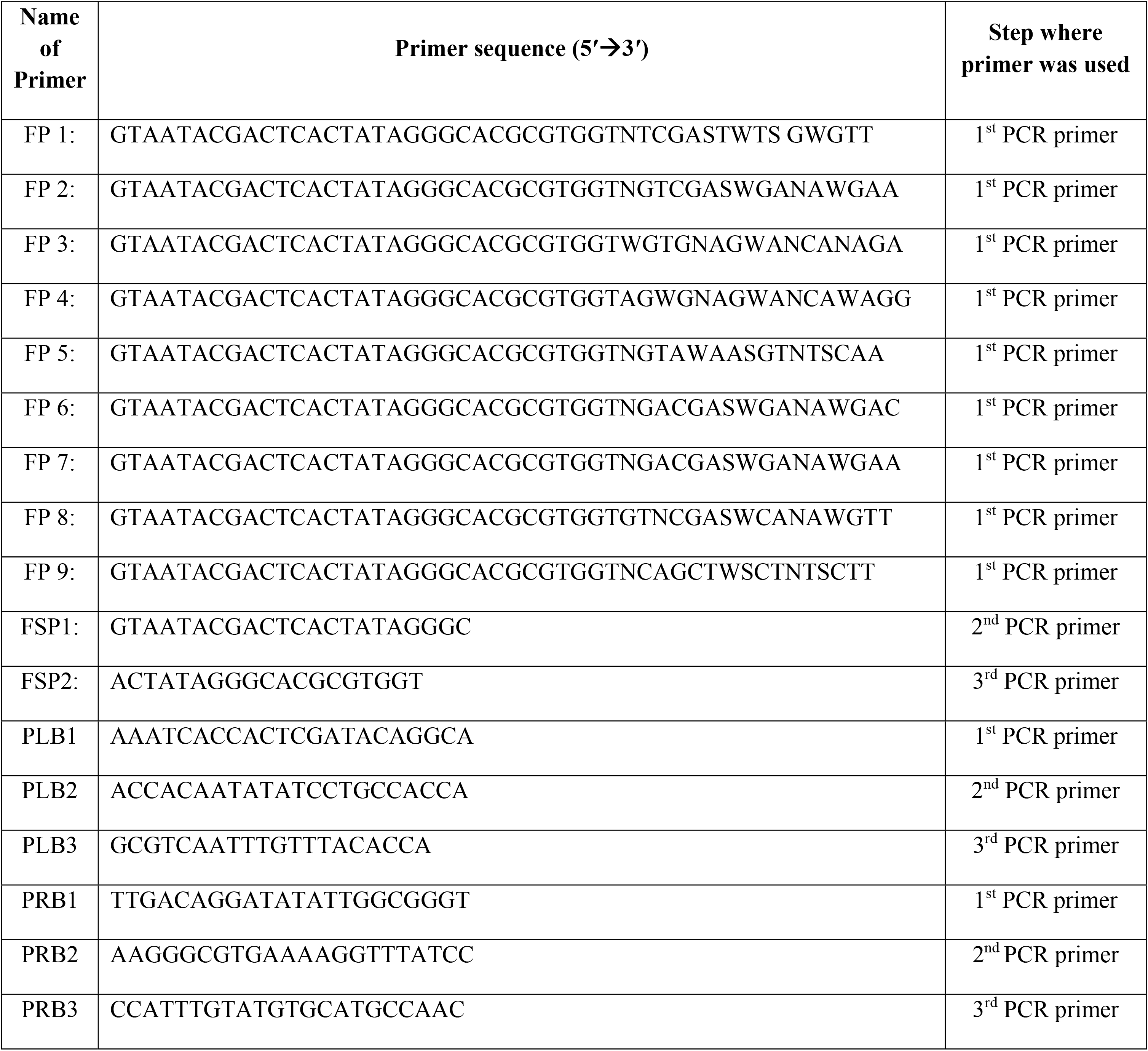
Sequences of primers used in FPNI-PCR. The PCR step at which the primers were used is mentioned at the right side of the respective primer.

**Table S7:**
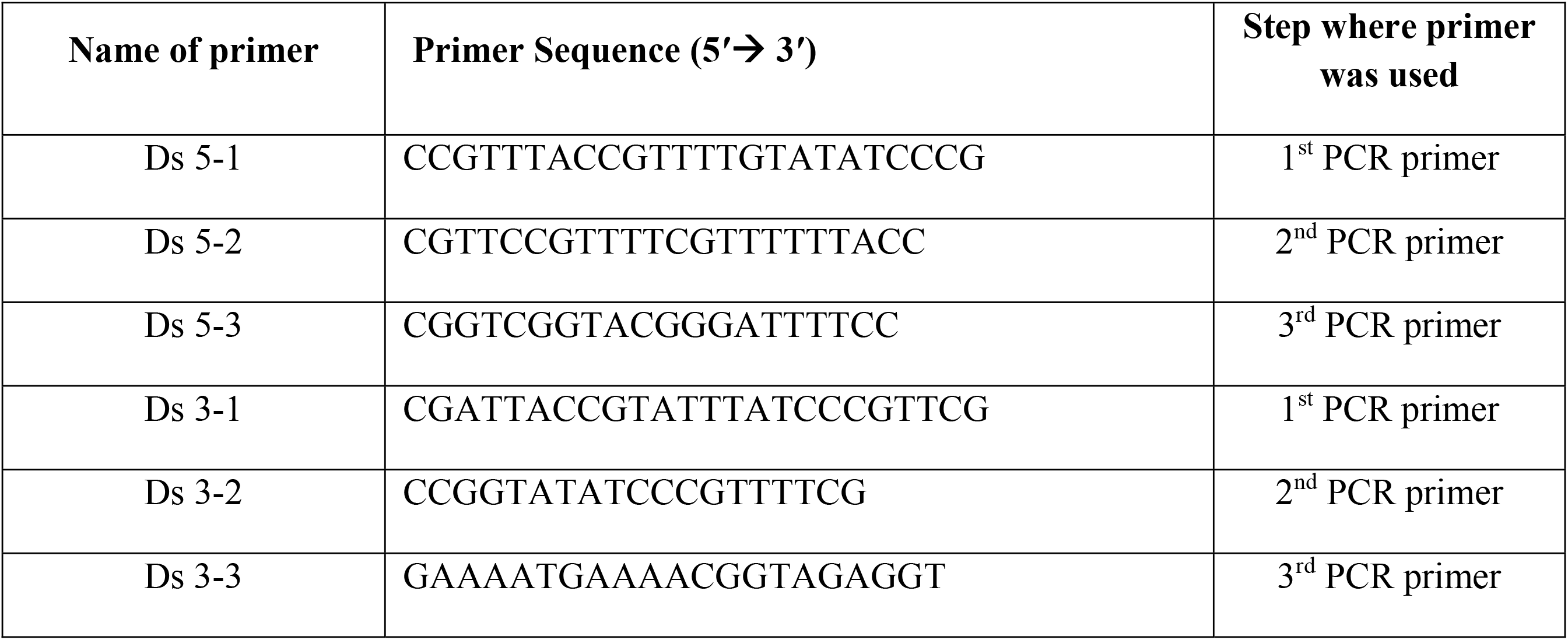
Sequences of primers used for inverse PCR. The step at which the primers were used is mentioned at the right side of the respective primer.

**Dataset S1:**
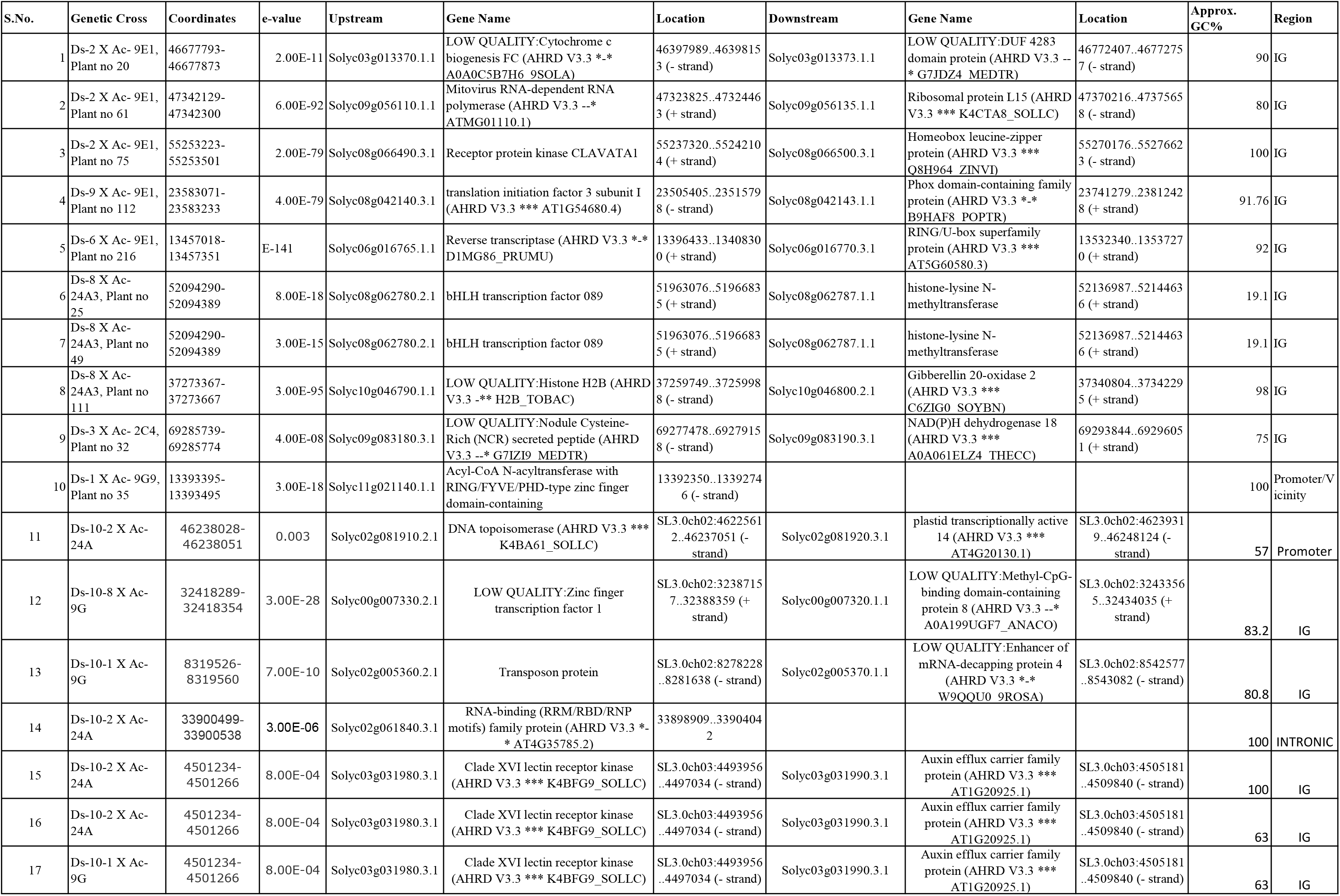
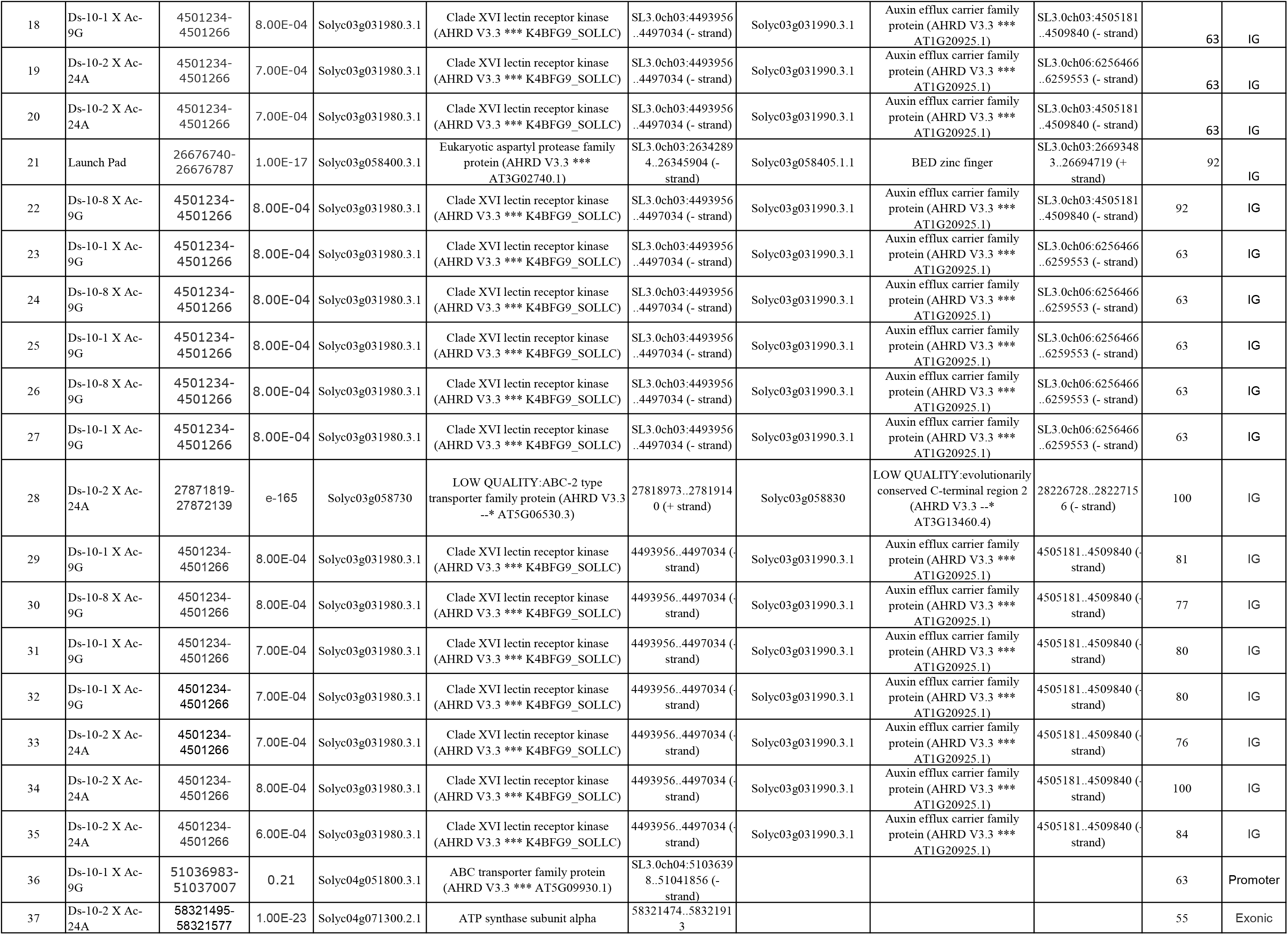
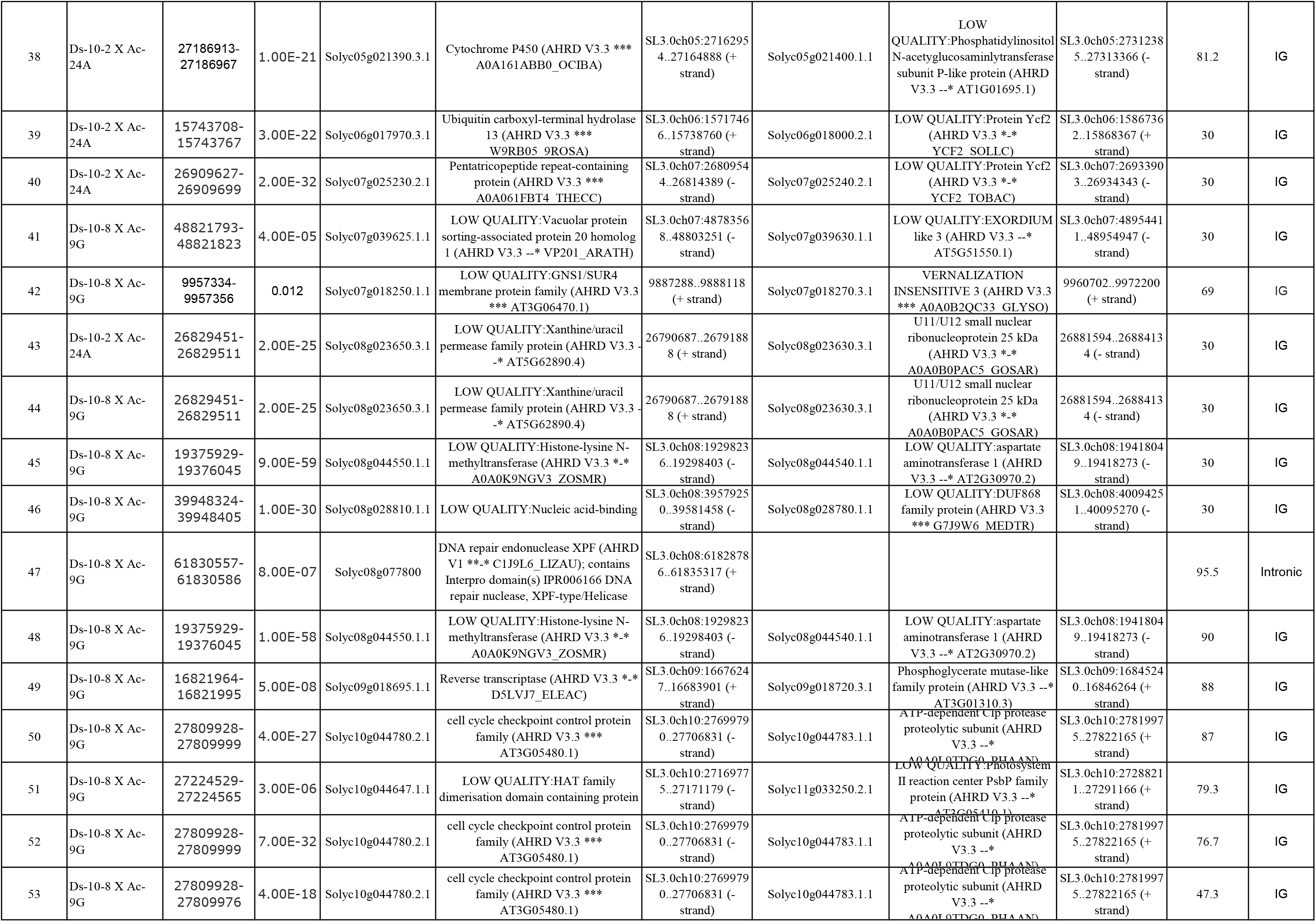
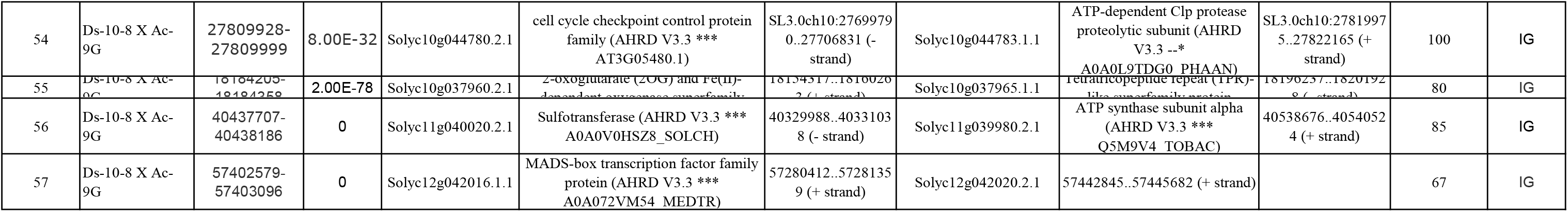
Location of *Ds* transposition in the tomato genome. The table shows the chromosomal coordinates of *Ds* transposition in tomato along with the e-value of the matching search. The names of genes upstream and downstream of transposed *Ds* elements are also indicated. The genic composition of transposed location is also mentioned.

